# *Salmonella* Typhimurium employ spermidine to exert protection against ROS-mediated cytotoxicity and rewires host polyamine metabolism to ameliorate its survival in macrophages

**DOI:** 10.1101/2023.09.29.560257

**Authors:** Abhilash Vijay Nair, Anmol Singh, R. S. Rajamani, Dipshikha Chakravortty

**Author notes:** Address correspondence to Dipshikha Chakravortty.

## Abstract

*Salmonella* infection involves a cascade of attacks and defence measures. After breaching the intestinal epithelial barrier, *Salmonella* is phagocytosed by the macrophages, inside which, the bacteria face multiple stresses and, consequently, employ appropriate countermeasures. We show that, in *Salmonella*, the polyamine spermidine activates a stress response mechanism by regulating critical antioxidant genes. *Salmonella* Typhimurium mutants for spermidine transport and synthesis cannot mount an antioxidative response, resulting in high intracellular ROS levels. These mutants are also compromised in their ability to be phagocytosed by macrophages. Furthermore, it regulates a novel enzyme in *Salmonella*, Glutathionyl-spermidine synthetase (GspSA), which is known to prevent the oxidation of proteins in *E.coli*. Moreover, the spermidine mutants and the GspSA mutant show significantly reduced survival in the presence of hydrogen peroxide *in vitro*, and lesser organ burden in the mouse model of *Salmonella* infection. Conversely, in macrophages isolated from *gp91phox*^-/-^ mice, we observed a rescue in the attenuated fold proliferation previously observed upon infection. Interestingly, *Salmonella* upregulates polyamine biosynthesis in the host through its effectors from SPI-1 and SPI-2, which also solves the mystery of the attenuated proliferation observed in spermidine transport mutants. Thus, inhibition of this pathway in the host abrogates the proliferation of *Salmonella* Typhimurium in macrophages. From a therapeutic perspective, inhibiting host polyamine biosynthesis using an FDA-approved chemopreventive drug, D,L-α-difluoromethylornithine (DFMO), reduces *Salmonella* colonization and tissue damage in the mouse model of infection, while enhancing the survival of infected mice. Therefore, our work provides a mechanistic insight into the critical role of spermidine in stress resistance of *Salmonella*. It also reveals a strategy of the bacteria in modulating host metabolism to promote their intracellular survival and shows the potential of DFMO to curb S*almonella* infection.

## Introduction

A condition with ample nutrients, and desired temperature, pH, oxygen concentration, and osmolarity is usually considered optimal for the growth of microbes. An imbalance or any alteration in these parameters impedes the growth and survival of the microbes, considered as stress conditions. However, these parameters keep fluctuating in nature [1]. Hence, to persist and survive in the natural environment, with unforeseen stressful or lethal conditions, they must be adept in sensing, responding and adapting to them [2]. In the case of food-borne pathogens, they face an array of various stresses in multiple environments in which they dwell, such as from the natural habitat to commercial settings and inside the host system [3–5]. *Salmonella* is a food-borne pathogen that enters the host system through contaminated food and water. In the host system, as it traverses to the intestine, *Salmonella* faces multiple stressful conditions such as low pH, nutrient deprivation, bile salt stress, competition with the resident microbes of the gut and immunoglobulins, etc . Once it breaches the epithelial barrier, it is taken up by the macrophages at the lamina propria, through which it disseminates throughout the host system. Macrophages are phagocytic immune cells where *Salmonella* encounters a very hostile environment. Entry of the pathogen into the cell cytoplasm leads to a burst of reactive oxygen species (ROS) and reactive nitrogen species (RNS) [6, 7]. In macrophages, *Salmonella* resides in a specialised niche called the *Salmonella*-containing vacuole (SCV), which presents multiple other stresses to the bacteria, such as acidification, nutrient limitation and attack by the antimicrobial peptides. However, *Salmonella* employs numerous weapons from its arsenal to counteract the stresses it faces within the host macrophages.

Polyamines are a group of primordial stress response molecules in prokaryotes and eukaryotes [8]. Multiple research groups have elucidated the link between polyamines and bacterial stress response. In *Shigella spp.* the polyamine spermidine accumulates during its infection into macrophages, which increases the expression of *katG* and helps in bacterial antioxidant response [9]. Moreover, spermidine localises to the surface of *Pseudomonas aeruginosa* and bind to the lipopolysaccharide (LPS) to protect the cells from oxidative damage [10]. In the Gram-positive bacteria, *Streptococcus pyogenes,* extracellular spermidine enhance the survival of the bacteria by upregulating oxidative response genes [11]. A research group has shown that polyamines are vital in resistance against nitrosative stress in *Salmonella* Typhimurium. Further, the group showed that spermidine is required for the systemic infection of *Salmonella* Typhimurium in mice [12]. Thus, it is conceivable that *Salmonella* Typhimurium utilises polyamines, such as spermidine, as a stress response molecule; however, the mechanism remains elusive.

Here, we show that spermidine biosynthesis and transport mutants of *Salmonella* Typhimurium exhibit reduced survival upon infection in RAW264.7 cells. This diminished proliferation is also observed in mice models of *Salmonella* infection, which is rescued in *gp91phox*^-/-^ mice. We demonstrate that spermidine orchestrates the various arms of antioxidative response and aids in tightly regulating intracellular ROS levels. We further identify a novel antioxidative enzyme, Gsp, in *Salmonella* Typhimurium, which is regulated by spermidine. The fascinating question that arises is why the transporter mutant shows reduced survival. To this, we find that *Salmonella* Typhimurium harnesses the host polyamine biosynthesis for its survival. Furthermore, for the first time, we show that an FDA-approved chemopreventive and anti-African Human Trypanosomiasis drug that inhibit the polyamine biosynthesis in the host, is able to curb *Salmonella* infection in mice models.

## Material and methods

### Bacterial strains and growth condition

*Salmonella enterica* serovars Typhimurium (STM WT) wild type strain ATCC 14028s was used in all experiments which was a kind gift from Prof. Michael Hensel, Abteilung Mikrobiologie, Universität Osnabrück, 273 Osnabrück, Germany. The bacterial strain was cultured in Luria broth (LB-Himedia) with constant shaking (175rpm) at 37°C orbital-shaker. Kanamycin, Chloramphenicol and Ampicillin antibiotics (Kanamycin-50μg/ml, Cholramphenicol-20μg/ml and Ampicillin-50μg/ml) were used wherever required. Strains were transformed with pFPV-m-cherry plasmid for immunofluorescence assays. (Bacterial strain list in Supplementary table **S-Table 1**)

### Bacterial gene knock-out and strain generation

The generation of gene knock-out in bacteria was done using the One-step chromosomal gene inactivation protocol [13]. Briefly, the Kanamycin resistance gene and the Chloramphenicol resistance gene amplified products were purified using chloroform-ethanol precipitation. Followed by electroporation into the STM WT cells (expressing PKD-46 plasmid which provides the λ-Red recombinase system) by a single pulse of 2.25 kV separately for the Kan^R^ and Chlm^R^. The transformant colonies were selected and patched on fresh plates and confirmed for knock-out using PCR with primers designed corresponding to the ∼100bp upstream and downstream of the genes (knocked out) for the knock-out strains to observe a difference in PCR product size upon STM *ΔpotCD* and STM *ΔspeED* knockout generation. For the generation of the double knock-out strain (STM *ΔpotCDΔspeED*), the STM *ΔpotCD* (resistant to Kanamycin) was first transformed with the plasmid pKD46 which provides the λ-Red recombinase system. To this transformed strain, the purified PCR product to knock-out *speED* was electroporated to generate the STM *ΔpotCDΔspeED* (resistant to Kanamycin and Chloramphenicol). For the generation of STM *Δgsp,* Kanamycin resistance gene was amplified from pKD4 plasmid, and a similar protocol was used, followed by selection on Kanamycin containing LB agar plates. For the generation of double gsp spermidine mutants, STM *Δgsp* was electroporated with purified PCR product to knock-out *speED* and *potCD* (both Chloramphenicol resistance cassette). (Knockout generation Primer list in Supplementary table **S-Table 2**)

### Cell culture and maintenance

RAW264.7 cells (murine macrophage cell line) were cultured in DMEM - Dulbecco’s Modified Eagle Medium (Lonza) supplemented with 10% FBS (Gibco) and 1% Penicillin-streptomycin (Sigma-Aldrich) at 37°C humidified chamber (Panasonic) with 5% CO_2_. For each experiment, the cells were seeded onto the appropriate treated cell culture well plate at a confluency of 80% for intracellular survival assay, and expression studies.

### Gentamicin protection assay

The cells were infected with STM WT, STM *ΔpotCD,* STM *ΔspeED,* STM *ΔpotCDΔspeED,* STM *Δgsp* and STM *ΔkatG* at MOI of 10 (for intracellular survival assay) and MOI 25 (for qRT-PCR). After infecting the cell line with STM WT and the mutants, the plate was centrifuged at 700-900 rpm for 10 minutes to facilitate the proper adhesion. The plate was then incubated for 25 minutes at 37°C humidified chamber and 5% CO_2_. Then the media was removed from the wells and washed with 1X PBS. Fresh media containing 100 µg/mL gentamicin was added and again incubated for 60 minutes at 37°C and 5% CO_2_. The media was then removed, cells were washed with 1X PBS twice, and fresh media containing 25µg/mL gentamicin was added. The plate was incubated at 37°C and 5% CO_2_ till the appropriate time. For the intracellular survival assay, two time points were considered 2 hours and 16 hours, and for qRT-PCR three time points were considered 2 hours, 6 hours and 16 hours. For phagocytosis assay upon opsonisation, all the strains were washed with 1XPBS and incubated at 37°C with mouse complement sera for 1 hour before gentamicin protection assay

### Intracellular survival assay and phagocytosis assay

At the appropriate time post-infection, the cells were lysed using 0.1% Triton X followed by addition of more 1X PBS and samples were collected. The collected samples were plated at the required dilutions on LB agar plates and kept at 37°C. 12 hours post incubation the Colony forming units (CFU) were enumerated for each plate.

The fold proliferation and invasion were determined as follows

Fold Proliferation = (CFU at 16 hours post-infection)/(CFU at 2 hours post-infection) Percentage Phagocytosis = [(CFU at 2 hours post-infection)/(CFU of the Pre-inoculum)]×100

### RNA isolation and qRT-PCR

RNA isolation was performed from infected cells after appropriate hours of infection with STM WT, STM *ΔpotCD,* STM *ΔspeED* by RNA isolation was performed using TRIzol (Takara) reagent according to manufactures’ protocol RNA was quantified using Thermo-fischer scientific Nano Drop followed by running on 2% agarose gel for checking the quality. For cDNA synthesis, first DNase I treatment with 3μg of isolated RNA was done at 37℃ for 60 minutes, which was then stopped by heating at 65℃ for 10 minutes. Then RNA (free of DNA) was subjected to Reverse transcription using Random hexamer, 5X RT buffer, RT enzyme, dNTPs and DEPC treated water at 37°C for 15 minutes, followed by heating at 85℃ for 15 seconds. Quantitative real-time PCR was done using SYBR green RT-PCR kit in BioRad qRT-PCR system. A 384 well plate with three replicates for each sample was used. The expression levels of the gene of interest were measured using specific RT primers. Gene expression levels were normalised to 16SrDNA primers of *S*. Typhimurium. Gene expression levels of eukaryotic gene of interest were normalised to beta-actin of mouse/human as required. For expression studies in bacteria grown in LB media, the bacterial samples were harvested at 3 hours, 6 hours, 9 hours and 12 hours post subculture in fresh LB media in 1:100 ratio and 1mM H_2_O_2_ was added to the broth, to study the gene expression in the presence of oxidative stress. Then similar protocol was used to isolate total RNA using TRIzol (Takara) reagent according to manufactures’ protocol (Expression Primer list in Supplementary table **S-Table 2**)

### Primary macrophages isolation and infection

Primary macrophages were isolated from C57BL/6 mice (male, 5-6 weeks old). The mice were intraperitoneally injected with 8% Brewer’s Thioglycolate (HiMedia). Five days post injection the primary macrophages were aseptically isolated by injecting ice-cold 1xPBS into the peritoneal cavity the peritoneal lavage was collected. Any residual erythrocytes were lysed using RBC lysis buffer (Sigma-R7757), and the isolated cells were maintained in complete RPMI 1640 media for further experiments.

### Intracellular Reactive oxygen species determination using H_2_DCFDA staining

Overnight cultures were sub-cultured in fresh LB media. Once the cultures reached OD 0.1 then 10^8^ CFU/ml of each strain was incubated with 10µM of 2’,7’-dichlorodihydrofluorescein diacetate (H2DCFDA) (Sigma) in 1xPBS (pH 7.2) at 37°C for 30 minutes. The bacterial cells were centrifuged and the cells were resuspended in 1xPBS (pH 7.2) with Hydrogen peroxide of different concentrations (0mM-10mM), and incubated 37°C (orbital shaker) for 2 hours. The samples were transferred to a 96 well ELISA plate and fluorescence was determined in Tecan-ELISA plate reader Infinite series 200 (Ex-490nm/ Em-520nm).

### Intracellular redox status determination

The STM WT, STM *ΔpotCD,* STM *ΔspeED* and STM *Δgsp* were transformed with pQE60-Grx1-roGFP2 (a gift from Prof. Amit Singh, CIDR, IISc). Overnight cultures were sub-cultured in fresh LB media. Once the cultures reached OD 0.1 then 10^8^ CFU/ml of each strain was incubated with Hydrogen peroxide (0mM-5mM) of different concentrations in 1xPBS (pH 7.2) with, and incubated 37°C (orbital shaker) for 2 hours. The samples were centrifuged and resuspended in fresh 1xPBS (pH 7.2). And tubes were analysed for GFP fluorescence at 408nm and 488nm respectively using BD-FACS Verse flow cytometer (total 10,000 events for each sample). For determination of intracellular redox status upon infection into RAW264.7 cells, 10^5^ RAW264.7 cells seeded in a 24 well tissue culture plate, were infected with each of the strains harboring the roGFP2 plasmid using gentamicin protection assay. After 16 hours post infection, the macrophages were washed with 1X PBS and scrapped off using cell scraper and analysed for GFP fluorescence at 408nm and 488nm respectively using BD-FACS Verse flow cytometer (total 10,000 events for each sample).

### *In vitro* sensitivity assays

Overnight cultures were sub-cultured in fresh LB media. Once the cultures reached OD 0.1 then 10^8^ CFU/ml of each strain was incubated with Hydrogen peroxide or sodium nitrite of different concentrations (0mM-10mM) in 1xPBS(pH 7.2) (1xPBS of pH 5.4 was used for nitrite sensitivity assay) with, and incubated 37°C (orbital shaker) for 2 hours. The samples were plated on SS agar to enumerate the CFU and the percentage survival was determined as:

Percentage Survival : [[CFU/ml for treated with H_2_O_2_/ NaNO_2_]/ [CFU/ml for untreated]]×100 Resasurin assay was used to determine the viable cells was performed in 96 well plate (triplicate for each sample and concentration. Briefly, after incubation for 2 hours as previously performed, Resasurin (0.2mg/ml in 1X PBS) was added (1:10 ratio) to each well of 96 well plate. The plate was incubated at 37°C shaker incubator for 2 hours. The fluorescence was measure using Tecan plate reader infinite series 200, with Ex-520nm and Em-590nm. The values obtained as Relative fluorescence units (RFU) and the percentage survival was determined as

Percentage survival = [RFU of sample with added hydrogen peroxide]/ [RFU of sample without of hydrogen peroxide]

### Immunoblotting

The bacterial strains were grown in LB media with added 1mM hydrogen peroxide until the log phase of growth. The cells were centrifuged to remove the media, and the cells were resuspended in the lysis buffer (Sodium chloride, Tris, EDTA, 10% protease inhibitor cocktail) after washing with 1XPBS. The cells were lysed using sonication and centrifuged at 4°C to collect the cell lysate, followed by estimation of total protein using the Bradford protein estimation method. 50µg of protein was loaded onto a Polyacrylamide Gel Electrophoresis (PAGE) without β-mercaptoethanol (prevent di-sulphde bond breakage as glutathionyl-spermidine modifies Cysteine residues through a disulphide bond), then transferred onto 0.45μm PVDF membrane (GE Healthcare). 5% skimmed milk (Hi-Media) in TTBS was used to block for 1hour at room temperature and then probed with Anti-Spermidine primary (Novus Biologicals) and the secondary HRP-conjugated antibodies. ECL (Biorad) was used for developing the blot, and images were captured using Chemi-Doc(Biorad). All densitometric analysis was performed using the Image J. The normalisation was done with respect to Ponceau S stained blot.

### Transfection for knockdown

RAW 264.7 cells were seeded at a confluency of 50-60% 12 hours prior to transfecting using either PEI (1:2 -DNA: PEI). We used two different constructs for knock-down of Odc1, A7 and F8 and both as a mixed construct (A7:A8 at 1:1 ratio) and similarly for Srm, E9 and F1 and both as a mixed construct (E9:F1 at 1:1 ratio) from the Sigma Mission shRNA library. 400ng of plasmid DNA/well (ratio 260/280 ∼1.8-1.9) was used for transfection in a 24well plate. Cells were then incubated for 8 hours at 37℃ in a humidified incubator with 5% CO2; after that, the media containing transfecting DNA and reagents were removed, and cells were further incubated for 48 hours in complete media DMEM +10% FBS. Cells were harvested for further analysis or infected with the required MOI using the gentamicin protection assay. (shRNA sequence list in Supplementary table **S-Table 3**)

### Immunofluorescence

After the appropriate incubation time after gentamicin protection assay, the media was removed, and the cells were washed with 1X PBS and fixed with 3.5% Paraformaldehyde for 10 minutes. The cells were then washed with 1X PBS, followed by incubation with the required primary antibodies (anti-mouseLAMP1 and anti-Spermidine) in a buffer containing 0.01% saponin and 2% BSA, and incubated at room temperature for 45-60 minutes. After washing with 1X PBS, the secondary antibody tagged to a fluorochrome was added and incubated (anti-rat-Alexafluor488 for LAMP1, anti-rabbit-Alexafluor647 for spermidine). The coverslips were then washed with PBS and mounted on a clean glass slide using mounting media containing an anti-fade reagent and observed under the confocal microscope (Zeiss 710 microscope, at 63X oil immersion, 2×319 3× zoom, and 100X zoom for studying only bacterial samples, Zeiss 880 microscope, at 63X oil immersion, 2×319 3× zoom).

The histopathological sections were deparaffinized and then incubated with the required primary antibody (anti-*Salmonella* LPS) in a buffer containing 0.01% saponin and 2% BSA and incubated at room temperature for 45-60 minutes. The primary antibody was then removed by washing with 1X PBS and then incubated with the appropriate secondary antibody tagged to a fluorochrome. The sections were then washed with PBS and cover with coverslip after using mounting media containing an anti-fade reagent. The coverslips were sealed with clear nail polish and observed under the confocal microscope. For studying histopathology samples Zeiss 880 microscope 40X oil immersion 2×319 3× zoom was used.

### Intracellular RNS determination

For determination of intracellular RNS, a cell permeable nitric-oxide probe, 4, 5-diaminofluorescein diacetate (DAF_2_A) (Sigma) was used. The protocol has been followed as Roy Chowdhury A et. al. [14]. Briefly, After 16 hours of infection of RAW264.7 cells (transiently knock down of *Odc1*) at MOI 10 with STM WT, the cells were incubated with fresh DMEM containing 5µM of DAF_2_DA. The cells were incubated at 37℃ in a humidified incubator with 5% CO_2_ for 30 minutes. The media containing dye was removed and the cells were washed with 1XPBS and the cells were acquired immediately for flow-cytometry (BD FACS Verse) (Ex-491nm/Em-513nm).

### Intracellular Glutathione determination

The intracellular reduced Glutathione (GSH) concentration was determined by modification of a pubmished protocol [15]. Briefly, a standard curve with known concentration of GSH (Sigma) was prepared. Reaction mixture for each contained 600µL of phosphate buffer (0.1M, pH7), 40µL of 0.4% w/v 5,5-dithiobis(2-nitrobenzoic acid) (DTNB, from Sigma), 100 µL of the standard solutions of GSH (0mM-1mM range) and autoclaved MilliQ water to make up the volume to 1000µL.The mixture was incubated at room temperature for 5 minutes and absorbance was measured at 412nm using tecan plate reader. The bacterial strains were subcultured in fresh LB media and grown till OD 0.1 (exponential phase) and washed with buffer (Tris, Sodium chloride and EDTA) and lysed using sonication. The supernatant after sonication was used as the sample for GSH detection. As previously described for standard curve. From the standard curve the intracellular concentrations were interpolated.

### In silico analysis

The *in silico* protein structure determination was performed using the SWISS-MODEL software (https://swissmodel.expasy.org/), where we supplied the protein sequence of GspSA of *Salmonella* Typhimurium from UniProt. We analysed the model with highest sequence identity and maximum coverage with the Gsp from *E.coli*. The structure depicts a Homo-dimer with a GMQE of 0.93 and QMEANisCo Global of 0.88 ± 0.05.

### *In vivo* animal experiment

5-6weeks old C57BL/6 mice were infected by orally gavaging 10^7^ CFU of STM WT, STM *ΔpotCD,* STM *ΔspeED,* STM *ΔpotCDΔspeED,* STM *Δgsp* and STM *ΔkatG*. To study the colonisation in organs, the intestine (Peyer’s patches), MLN, spleen and liver were isolated aseptically, 5 days post-infection, and the CFU was enumerated on differential and selective SS agar by serial dilution. For intraperitoneal infection 5-6weeks old C57BL/6 mice were infected by intraperitoneally injecting 10^3^ CFU of STM WT, STM *ΔpotCD,* STM *ΔspeED,* STM *ΔpotCDΔspeED,* STM *Δgsp* and STM *ΔkatG.* To study the colonisation in organs, spleen and liver were isolated aseptically 3 days post-infection. Blood was isolated by heart puncture 3 days post-infection. The CFU was enumerated on differential and selective SS agar by serial dilution. Organs were stored in 3.5%PFA before histopathogical sample preparation.

For inhibitor treatment, 5-6weeks old C57BL/6 mice were infected by orally gavaging 10^7^ CFU of STM WT. The inhibitor DFMO (Sigma D193) was intraperitoneally injected every alternate day from day 1 at two doses 2mg/kg and 1mg/kg of body weight of mice. To study the colonisation in organs, the intestine (Peyer’s patches), MLN, spleen and liver were isolated aseptically, 5 days post-infection, and the CFU was enumerated on differential and selective SS agar by serial dilution. For survival assay of mice 5-6weeks old C57BL/6 mice were infected by orally gavaging 10^8^ CFU of STM WT. The inhibitor DFMO was intraperitoneally injected every alternate day from day 1 at two doses 2mg/kg and 1mg/kg of body weight of mice. The survival was monitored till 10 days. Likewise, organs were stored in 3.5% PFA before histopathogical sample preparation.

### Mass Spectrometry to determine intracellular GS-sp levels

The bacterial strains were grown in LB media with added 1mM hydrogen peroxide until the log phase of growth. The cells were centrifuged to remove the media, and the cells were resuspended in the lysis buffer (Sodium chloride, Tris, EDTA, 10% protease inhibitor cocktail) after washing with 1XPBS. The cells were lysed using sonication and centrifuged at 4°C to collect the cell lysate. Protein was precipitated using ice-cold acetone (Sigma, MS grade), 4 times the volume of the cell lysate and by incubating at -20°C overnight. The precipitated proteins were removed by centrifugation and the supernatatnt was used for analysis. Samples were analysed by ESI MS Q-TOF, impact HD (Bruker Daltonics Germany) connected to Agilent HPLC 1260. Samples were passed through Agilent C18, 4.6X150mm column. Mobile phase used was water and Acetonitrile with 0.1% formic acid. Linear gradient was used with flow rate 0.2ml/min. Data was analysed using Bruker Daltonics software Data analysis 4.1. Mass of GS-sp (Glutathionyl spermidine) is 434g/mol, and (GS-sp)_2_ (oxidized form, Di-glutathionyl spermidine) is 866g/mol.

### Statistical Analysis

Statistical analyses were performed with GraphPad Prism software. The Student’s t-test (parametric, two-tailed, unpaired) was performed as indicated. For animal experiments Non-parametric Mann-whitney (two-tailed) test was performed. The results are expressed as mean ± SD or mean ± SEM. Group sizes, experiment number, and p values for each experiment are described in figure legends.

## Results

### Loss of spermidine transporter and biosynthesis genes in *Salmonella* Typhimurium compromises its ability to be phagocytosed by the macrophages

The pathoadaptation in *Salmonella* involves multiple players, which counteracts the stressful condition it encounters in the host macrophages. Polyamines being a group of well-studied stress response molecules, we were interested in determining the expression of the spermidine transporter and biosynthesis genes in *Salmonella* Typhimurium. Our previous study shows that *Salmonella* upregulates the spermidine transporter genes (*potA, potB, potC* and *potD*) and the biosynthesis genes (*speE* and *speD*) during the log phase of growth *in vitro* [16]. Here we assessed the mRNA levels of *potA, potB, potC* and *potD* in STM WT upon infection into the RAW264.7 macrophage cell line. We noted that all the genes showed a 1.5-2 fold upregulation post 6 hours of infection into macrophages till 16 hours (**Fig 1 A)**. Further, our results showed that the spermidine biosynthesis enzymes *speE* and *speD* were upregulated 1.5 to 2 folds post 6 hours to 16 hours post-infection into macrophages (**Fig 1 B)**. These results indicate that *Salmonella* Typhimurium enhances its intracellular spermidine biosynthesis and imports from the extracellular milieu. It might be a strategy of the pathogen to increase the intracellular pool of stress response molecules like the polyamine spermidine, as it encounters the hostile environment of host macrophages.

**Figure 1:**
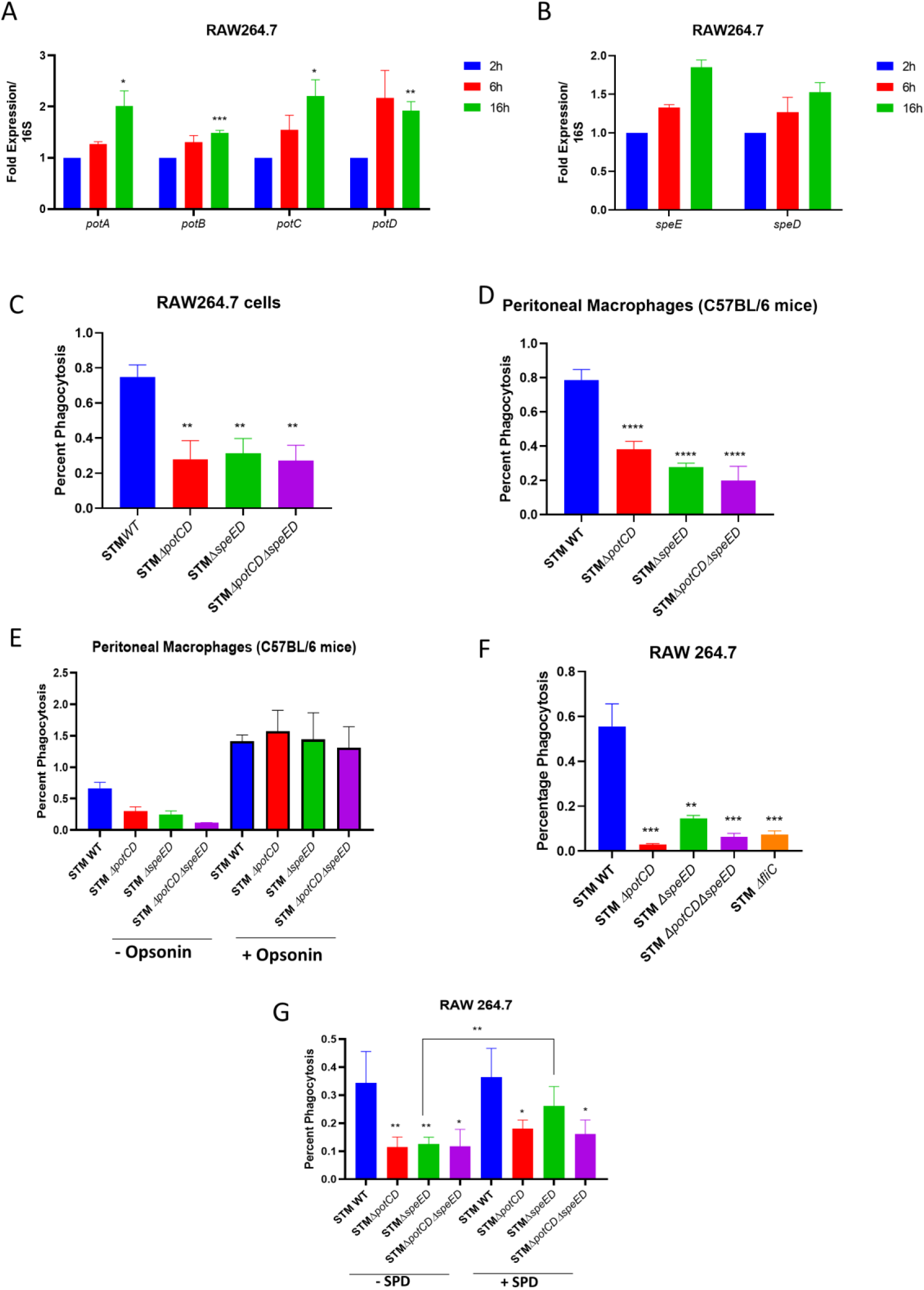
Loss of spermidine transporter and biosynthesis genes in *Salmonella* Typhimurium compromises its ability to be phagocytosed by the macrophages. A.The mRNA expression of spermidine transporter genes *potA, potB, potC* and *potD* in STM WT upon infection into RAW264.7 cells, B. The mRNA expression of spermidine biosynthesis genes *speE* and *speD* in STM WT upon infection into RAW264.7 cells, C. The percentage phagocytosis of the spermidine mutants in RAW264.7 cells, D. The percentage phagocytosis in primary macrophages isolated from peritoneal lavage of C57BL/6 mice, E. The percentage phagocytosis upon pre treatment with mouse-complement sera which act as an opsonin, F. The percentage phagocytosis in RAW264.7 cells with flagellin mutant (STM *ΔfliC*), G. The percentage phagocytosis in RAW264.7 cells with the spermidine mutants grown in media supplemented with 100µM spermidine (SPD). Student’s t-test was used to analyze the data; p values ****<0.0001, ***<0.001, **<0.01, *<0.05. Two-way Anova was used to analyze the grouped data; p values ****<0.0001, ***<0.001, **<0.01, *<0.05.

We then determined how the spermidine transporter and biosynthesis mutants (STM *ΔpotCD*, STM *ΔspeED* and STM *ΔpotCDΔspeED*) behave upon infection into RAW264.7 macrophages. We infected STM WT, STM *ΔpotCD*, STM *ΔspeED* and STM *ΔpotCDΔspeED* into RAW264.7 cells and observed that STM *ΔpotCD*, STM *ΔspeED* and STM *ΔpotCDΔspeED* showed reduced phagocytosis by the macrophages compared to the wild type (**Fig 1 C)**. Interestingly, STM *ΔpotCD*, STM *ΔspeED* and STM *ΔpotCDΔspeED* showed a compromised ability to proliferate intracellularly in the RAW264.7 cells further validating a previous study [12](**Fig S1 A)**. To further validate our results, we assessed the behaviour of the spermidine transporter and biosynthesis gene mutants in primary macrophages isolated from the peritoneal lavage of C57BL/6 mice. Likewise, the mutants showed attenuated proliferation and uptake by phagocytosis into the peritoneal macrophages (**Fig 1 D and S1 B)**. Collectively these results suggest that spermidine is a critical molecule in *Salmonella* Typhimurium to infect and survive in macrophages. To further investigate the reason behind the reduced ability to be taken up by macrophages upon loss of spermidine biosynthesis and transport, we treated STM WT, STM *ΔpotCD*, STM *ΔspeED* and STM *ΔpotCDΔspeED* with mouse complement-sera. Mouse complement-sera acts as an opsonin and thus potentiates the interaction of the bacteria with the macrophages. Upon pre-treatment with mouse complement-sera, we noted a rescue in the reduced uptake of the mutants by peritoneal macrophages isolated from C57BL/6 mice (**Fig 1 E)**. A study on *Salmonella* Typhimurium revealed that the aflagellate and non-motile *Salmonella* collide less frequently with macrophages and gets merest time to maintain contact with the macrophages, thereby showing decreased phagocytosis [17]. Our group previously showed that loss of spermidine production and import in *Salmonella* Typhimurium results in the loss of flagella formation on the bacterial cell surface [16]. Thus, we determined the percentage phagocytosis for a flagellin-deficient strain of *Salmonella* Typhimurium, STM *ΔfliC* and observed that STM *ΔpotCD*, STM *ΔspeED*, STM *ΔpotCDΔspeE* and STM *ΔfliC* exhibited a significantly decreased ability to be taken up by the RAW264.7 macrophages (**Fig 1 F**). Furthermore, we observed a rescue of the attenuated percentage phagocytosis and fold proliferation of only STM *ΔspeED* upon supplementation of spermidine (100µM) during the growth of bacteria prior to infection (**Fig 1 G and S1C**). We generated single gene mutants for abrogating the spermidine transport (STM *ΔpotA*) and spermidine biosynthesis (STM *ΔspeE*) function and further complemented the genes through a vector (STM *ΔpotA:potA* and STM *ΔspeE:speE*), we observed a recovery of the fold proliferation and percentage phagocytosis nearly to STM WT in the complemented strains (**Fig S1 D-E)** . Hence, the plausible explanation is that the reduced ability to form flagella in the spermidine mutants causes less frequent interaction with the macrophages and provides minimal contact time for infection, leading to reduced phagocytosis by the macrophages.

### Spermidine provides stress resistance in *Salmonella* Typhimurium by regulation of the expression of numerous antioxidative enzymes

The loss of spermidine transport and biosynthesis function in *Salmonella* Typhimurium renders it incapable of proliferation and survival in macrophages. In the host macrophages, the bacteria encounter numerous threats, of which the foremost is the rapid oxidative burst mediated by the NOX2. The reactive oxygen species superoxide radical can easily diffuse through the bacterial membrane and pose a major threat to the pathogen. ROS acts on multiple molecules such as nucleic acids, proteins and lipids, thus damaging the cell membranes, DNA and proteins within the bacteria [18]. Spermidine has been linked to stress response against oxidative stress and protects bacteria in *E. coli* and *Pseudomonas aeruginosa.* Thus, we were intrigued to understand the role of spermidine in antioxidative response in *Salmonella* Typhimurium. We examined the survival of STM WT, STM *ΔpotCD*, STM *ΔspeED*, STM *ΔpotCDΔspeE* upon exposure to the oxidative agent hydrogen peroxide *in vitro.* We noticed that at high concentrations of hydrogen peroxide, 5mM and 10mM, the STM *ΔpotCD*, STM *ΔspeED* and STM *ΔpotCDΔspeE* showed significantly lesser survival than STM WT(**Fig 2 A**). Moreover, the complemented strains for spermidine transport and synthesis mutants showed a rescue in attenuated survival in the presence of high concentrations of hydrogen peroxide *in vitro* (**Fig S2 A)**. We further determined the expression of *potA, potB, potC, potD*, *speE* and *speD* in STM WT upon exposure to 1mM hydrogen peroxide. There was a 2-3 fold upregulation in the mRNA expression of the transporter genes over the untreated during the early log phase of growth (6 hours) and the late log phase of growth (12 hours) *in vitro* (**Fig S2 B-E**). Similarly, the biosynthesis genes *speE* and *speD* were 4-6 fold upregulated in their corresponding mRNA expressions during their early log phase of growth (6 hours) and the late log phase of growth (12 hours) *in vitro* (**Fig S2 F and G**). Our results show that *Salmonella* Typhimurium upregulates spermidine transport and biosynthesis upon oxidative stress suggesting that spermidine mounts a protective function in such a stressful condition to aid bacterial survival.

**Figure 2:**
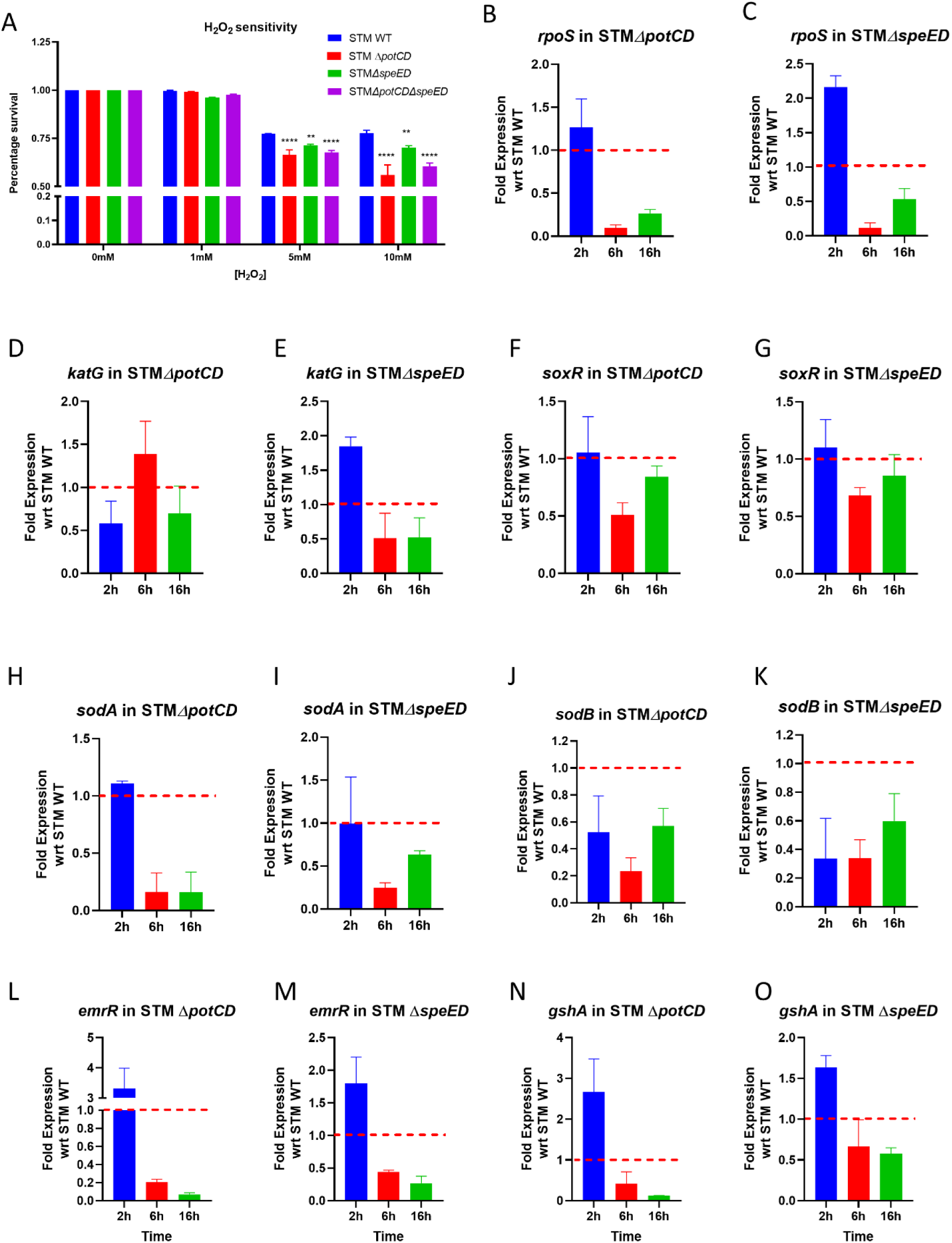
Spermidine provides stress resistance in *Salmonella* Typhimurium by regulation of the expression of numerous antioxidative enzymes. A.The *in vitro* hydrogen peroxide sensitivity assay with the spermidine transport and biosunthesis mutants, B. The mRNA expression of stress responsive transcription factor *rpoS* in STM *ΔpotCD* upon infection into RAW264.7 cells, C. The mRNA expression of stress responsive transcription factor *rpoS* in STM *ΔspeED* upon infection into RAW264.7 cells, D. The mRNA expression of *katG* in STM *ΔpotCD* upon infection into RAW264.7 cells, E. The mRNA expression of *katG* in STM *ΔspeED* upon infection into RAW264.7 cells, F. The mRNA expression of oxidative stress responsive transcription factor *soxR* in STM *ΔpotCD* upon infection into RAW264.7 cells, G. The mRNA expression of oxidative stress responsive transcription factor *soxR* in STM *ΔspeED* upon infection into RAW264.7 cells, H. The mRNA expression of *sodA* in STM *ΔpotCD* upon infection into RAW264.7 cells, I. The mRNA expression of *sodA* in STM *ΔspeED* upon infection into RAW264.7 cells, J. The mRNA expression of *sodB* in STM *ΔpotCD* upon infection into RAW264.7 cells, K. The mRNA expression of *sodB* in STM *ΔspeED* upon infection into RAW264.7 cells, L. The mRNA expression of glutathione synthetase specific transcription factor *emrR* in STM *ΔpotCD* upon infection into RAW264.7 cells, M. The mRNA expression of glutathione synthetase specific transcription factor *emrR* in STM *ΔspeED* upon infection into RAW264.7 cells, N. The mRNA expression of *gshA* in STM *ΔpotCD* upon infection into RAW264.7 cells, O. The mRNA expression of *gshA* in STM *ΔspeED* upon infection into RAW264.7 cells.

Bacteria sense the environmental changes and cues to respond and adapt to the altered environment. They use the two-component systems, transcriptional activators and repressors to alter gene expression in response to a stimulus [19]. Polyamines in *E. coli* regulate multiple genes at the transcription and translation together, referred to as the “Polyamine modulon”. These involve the numerous mRNAs, tRNAs, sigma factors, translational factors and two-component systems during the bacterial growth as well as in stress conditions [20–22]. *Salmonella* harbors multiple antioxidative enzymes to detoxify the ROS intracellularly [23]. Our study so far show that STM *ΔpotCD*, STM *ΔspeED* and STM *ΔpotCDΔspeE* is attenuated in survival under *in vitro* oxidative stress than STM WT. To gain mechanistic insight into the attenuated survival of *Salmonella* Typhimurium we determined the mRNA expression of the critical transcription factor *rpoS*, which activates the expression of the catalase enzymes (*katG and katE*) in order to detoxify hydrogen peroxide to water in the bacteria enzymatically [24]. In both STM *ΔpotCD* and STM *ΔspeED* the mRNA expression of *rpoS* is significantly down regulated from 6 hours post infection in RAW264.7 cells (**Fig 2 B and C, S3 A**). Further, its downstream target *katG* was also down regulated in STM *ΔpotCD* and STM *ΔspeED* from 6 hours post infection into RAW264.7 cells (**Fig 2 D and E, S3 B**). We further assessed the mRNA expression of the transcription factor *soxR*, which regulates the expression of superoxide dismutases (*sodA* and *sodB*) [25]. Upon infection in RAW264.7 cells, expression of *soxR* was significantly downregulated in STM *ΔpotCD* and STM *ΔspeED* with respect to STM WT (**Fig 2 F and G, S3 C**). Superoxide dismutases act on superoxide radicals, the potent ROS encountered in macrophages, converting to hydrogen peroxide. The mRNA expression of both *sodA* and *sodB* were likewise downregulated in STM *ΔpotCD* and STM *ΔspeED* upon infection into RAW264.7 macrophages (**Fig 2 H-K, S3 D and E**). A major antioxidant in most living organisms is glutathione (GSH), which directly acts as a quencher of ROS [26]. GSH is synthesized by Glutathione synthase (GshA) which in turn is regulated by EmrR transcription factor. We observed that the mRNA expression of *emrR* is downregulated in STM *ΔpotCD* and STM *ΔspeED* upon infection into RAW264.7 macrophages (**Fig 2 L and M, S3 F**). Similarly, *gshA* transcript expression is downregulated as well (**Fig 2 N and O, S3 G)**. To further validate the down regulation of the glutathione synthesis arm in spermidine transport and biosynthesis mutants, we determined the intracellular GSH levels and noted the in STM *ΔpotCD,* STM *ΔspeED* and STM *ΔpotCDΔspeED* the levels were significantly less (**Fig S3H and I).** Taken together these results indicate that spermidine regulates the transcription of multiple transcription factors involved in oxidative stress response in *Salmonella* Typhimurium. Importantly, we found a mechanism of oxidative stress resistance in *Salmonella* Typhimurium regulated by the spermidine, potentiating the survival of the bacteria.

### Spermidine controls a novel enzyme Glutathionyl-spermidine synthetase in *Salmonella* Typhimurium, and together mount an intracellular antioxidative response

The spermidine synthesized from putrescine has two fates. It is either acetylated by the enzyme SpeG or it is covalently conjugated to GSH to form Glutathionyl spermidine (GS-sp) catalysed by the enzyme Glutathionyl-spermidine synthetase (Gss). Tabor (1974) first discovered the existence of this enzyme in *E. coli* [27]. In *E. coli* GS-sp is generated at a higher level in the stationary phase and very less in the late exponential phase. It also interacts and modifies the thiol containing proteins under high H_2_O_2-_containing media leading to the formation of Gsp-thiolated proteins (PS-Gsp). Certain *in vitro* experiments with dehydro-ascorbate suggested that GS-sp might have higher antioxidant properties than GSH [28] and may be more effective in protecting against DNA damage by free radicals [29]. However, in *E. coli* Gss could not be linked to its pathogenicity [30]. Among *Enterobacteriaceae*, *Salmonella* was found to possess this unique enzyme. Our *in silico* analysis suggested that the enzyme in *Salmonella* Typhimurium (GspSA, encoded by *gsp*) has 90% identity with *E. coli gss* and the SWISS MODEL predicts it to be a homo-dimeric protein (**Fig S4 A-C**). Also, the spermidine synthesised by SpeE in *Salmonella* Typhimurium is directly fed into the pathway to synthesise GS-sp. We thus, investigated biological role of this novel enzyme in *Salmonella* Typhimurium. We noted that the mRNA expression of *gsp* is significantly upregulated at the late-log phase of growth of STM WT in LB media in the presence of H_2_O_2_ (**Fig S4 D**). Upon infection into RAW264.7 cells, STM WT upregulates the mRNA expression of *gsp* at 6 hours and 16 hours post-infection into RAW264.7 cells (**Fig 3 A**). Further, the *Salmonella* Typhimurium mutant of *gsp* (STM *Δgsp*) showed attenuated proliferation in RAW264.7 cells similar to STM *ΔkatG,* which has reduced capability to detoxify ROS (**Fig 3 B)**. Thus, our results suggest that *gsp* is important in *Salmonella* Typhimurium to survive and cope with the oxidative stress and hostile environment of macrophages. Interestingly, the mRNA expression of *gsp* was found to be downregulated in both STM *ΔpotCD* and STM *ΔspeED* at 16 hours post-infection into RAW264.7 macrophages (**Fig 3 C and D)**. These findings show that spermidine maintains its intracellular levels by regulating the flux of spermidine into other pathways, such as the GspSA pathway. Also, it further potentiates the ability of *Salmonella* Typhimurium to mount an antioxidative response by regulating *gsp* expression.

**Figure 3:**
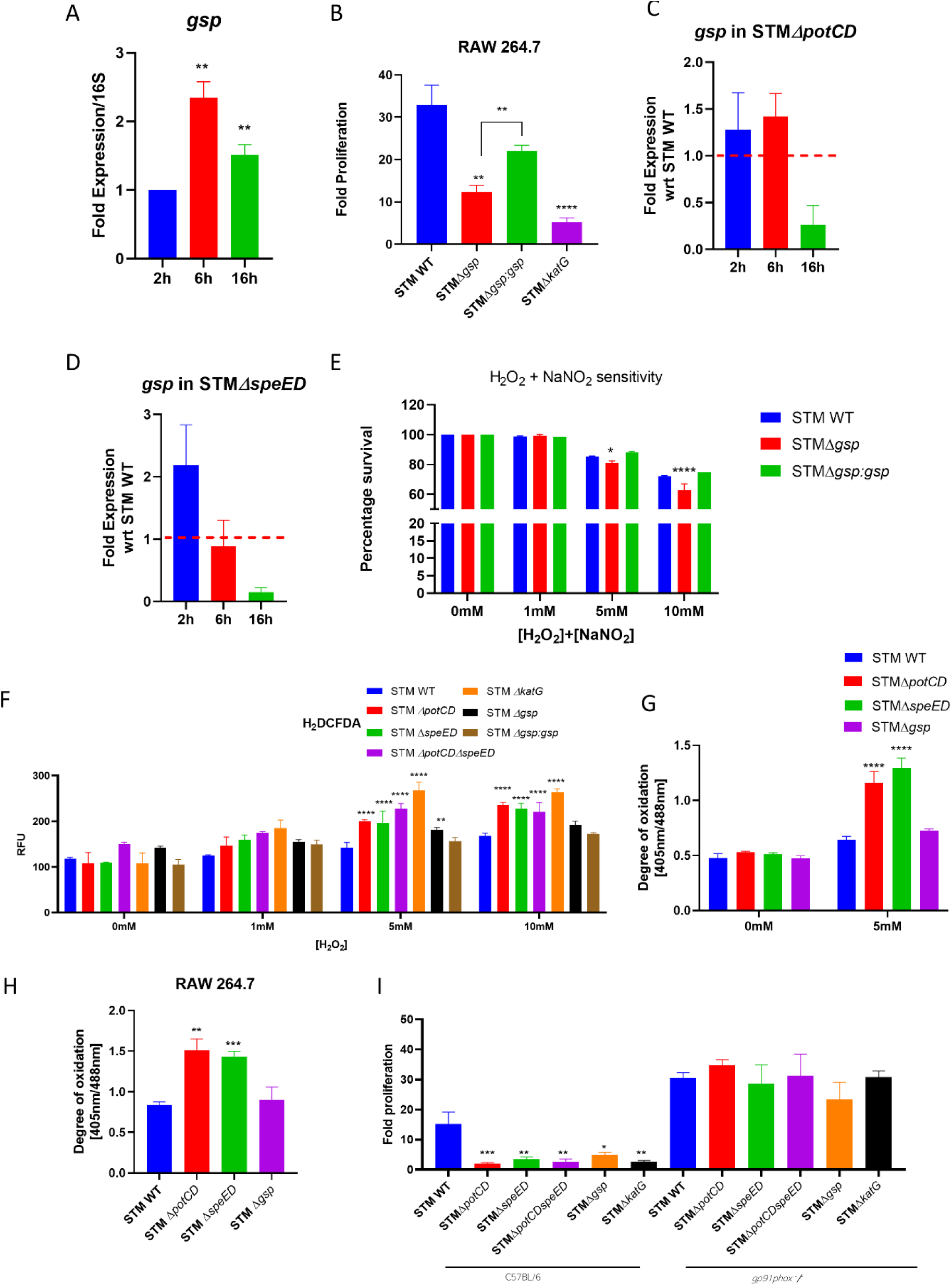
Spermidine controls a novel enzyme Glutathionyl-spermidine synthetase in *Salmonella* Typhimurium, and together mount an intracellular antioxidative response. A.The mRNA expression of *gsp* in STM WT upon infection into RAW264.7 cells, B The fold proliferation of STM WT, STM *Δgsp* and STM *Δgsp:gsp* in RAW264.7 cells, C. The mRNA expression of *gsp* in STM *ΔpotCD* upon infection into RAW264.7 cells, D. The mRNA expression of *gsp* in STM *ΔspeED* upon infection into RAW264.7 cells, E The *in vitro* Hydrogen peroxide and nitic oxide sensitivity assay, F. Intracellular reactive oxygen species determination using the cell permeable H_2_DCFDA dye, G. The intracellular degree of oxidation in spermidine and *gsp* mutants using pQE60-grx1-roGFP2 construct *in vitro*, H. The intracellular degree of oxidation using pQE60-grx1-roGFP2 construct in spermidine and *gsp* mutants upon infection into RAW 264.7 cells, I. The fold proliferation in primary macrophages isolated from wild type C57Bl/6 mice and *gp91phox-/-* mice, Student’s t-test was used to analyze the data; p values ****<0.0001, ***<0.001, **<0.01, *<0.05. Two-way Anova was used to analyze the grouped data; p values ****<0.0001, ***<0.001, **<0.01, *<0.05.

GspSA enzyme in Trypanosomes was named Trypanothione synthetase, while the conjugate is called Trypanothione (TSH). In trypanosomes TSH is essential, and these organisms entirely rely on TSH and possess no GSH reductase. Also, they have evolved to use TSH reductase instead of GSH reductase, Glutaredoxins or Thioredoxins [31]. In *E. coli* GS-sp forms bonds with the Cysteine thiol groups of numerous proteins and protect them from oxidation under oxidative stress. Cysteine thiol groups are highly prone to attack by ROS and get oxidized to sulphinic and sulphonic acids. In *Salmonella* Typhimurium we observe that loss of *gsp* results in attenuated proliferation and survival in macrophages. Thus, we investigated the survival of STM *Δgsp* upon exposure to hydrogen peroxide *in vitro*. STM *Δgsp* exhibited reduced survival in the presence of high 5mM and 10mM concentrations of H_2_O_2_ (**Fig S4 E**). Likewise, upon exposure to agents of oxidative stress and nitrosative stress H_2_O_2_ and NaNO_2_ together, we observe that STM *Δgsp* exhibited reduced survival at higher concentrations such as 5mM and 10mM (**Fig 4 E**). Thus, *gsp* is critical in *Salmonella* Typhimurium to shield the bacteria from the action of ROS and RNS. As we observed that spermidine transporter and biosynthesis mutants and *gsp* mutant of *Salmonella* Typhimurium are compromised in their survival under oxidative stress and in macrophages thus, we were interested to assess the intracellular ROS detoxification abilities of the strains. We determined the intracellular ROS in STM WT, STM *ΔpotCD*, STM *ΔspeED*, STM *ΔpotCDΔspeE,* STM *Δgsp* and STM *ΔkatG*. Interestingly, STM *ΔpotCD*, STM *ΔspeED*, STM *ΔpotCDΔspeE,* STM *Δgsp* and STM *ΔkatG* showed significantly higher intracellular ROS levels when they were exposed to 5mM and 10mM H_2_O_2_ (**Fig 3 F**). Further the complemented strains for spermidine transport and synthesis mutants showed a reduced intracellular ROS in the presence of high concentrations of hydrogen peroxide *in vitro* (**Fig S4 F)**. These results suggest that lower intracellular levels of spermidine correlate to higher intracellular ROS levels in *Salmonella* Typhimurium. To validate our observed results, we utilised a genetically engineered tool to sense the redox status of the bacterial cytosol, roGFP2 is a genetically modified form of GFP, and we use pQE60-Grx1-roGFP2 plasmid. The glutaredoxin (Grx1) fused to the roGFP2, reversibly transfers electrons between the cytosolic pool of GSH/GSSG and the thiol group of roGFP2, and the ratio of fluorescence ratio at 408nm and 488nm determine the redox status of bacterial cytoplasm [14]. We observed that STM *ΔpotCD*, STM *ΔspeED* harbouring the pQE60-Grx1-roGFP2 showed higher ratio of 405nm/488nm compared to STM WT in the presence of 5mM H_2_O_2_ *in vitro* and also upon infection into RAW 264.7 macrophages (**Fig 3 G and H**). However, STM *Δgsp* did not show a significantly higher ratio of 405nm/488nm. Moreover upon supplementation of the growth media with 100 µM spermidine, only in STM *ΔspeED* we observed a lower intracellular ROS and lesser 405nm/488nm in higher concentration of H_2_O_2_ (**Fig S4 G and H)**. Thus, our results indicate that spermidine is critical in mounting an antioxidative response to detoxify the intracellular ROS, by regulating multiple antioxidant genes in *Salmonella* Typhimurium. To validate our observed results we determined the intracellular levels of glutathionyl-spermidine in STM WT, STM *ΔpotCD*, STM *ΔspeED* and STM *Δgsp* using mass spectrometry. Our study qualitatively shows that the synthesis of GS-sp and (GS-sp)_2_ (oxidized form, di-glutathionylspermidine) in STM WT upon exposure to 1mM hydrogen peroxide, which is absent in the spermidine mutants and STM *Δgsp* (**Fig S5 A and B**). We further determined the presence of PS-Gsp and show that STM WT shows modifications of proteins by GS-sp (detected by anti-spermidine antibody), which is reduced upon treatment with beta-mercaptoethanol. The spermidine mutants show very less modification, while it is almost negligible in STM *Δgsp* (**Fig S5 C-E**).

**Figure 4:**
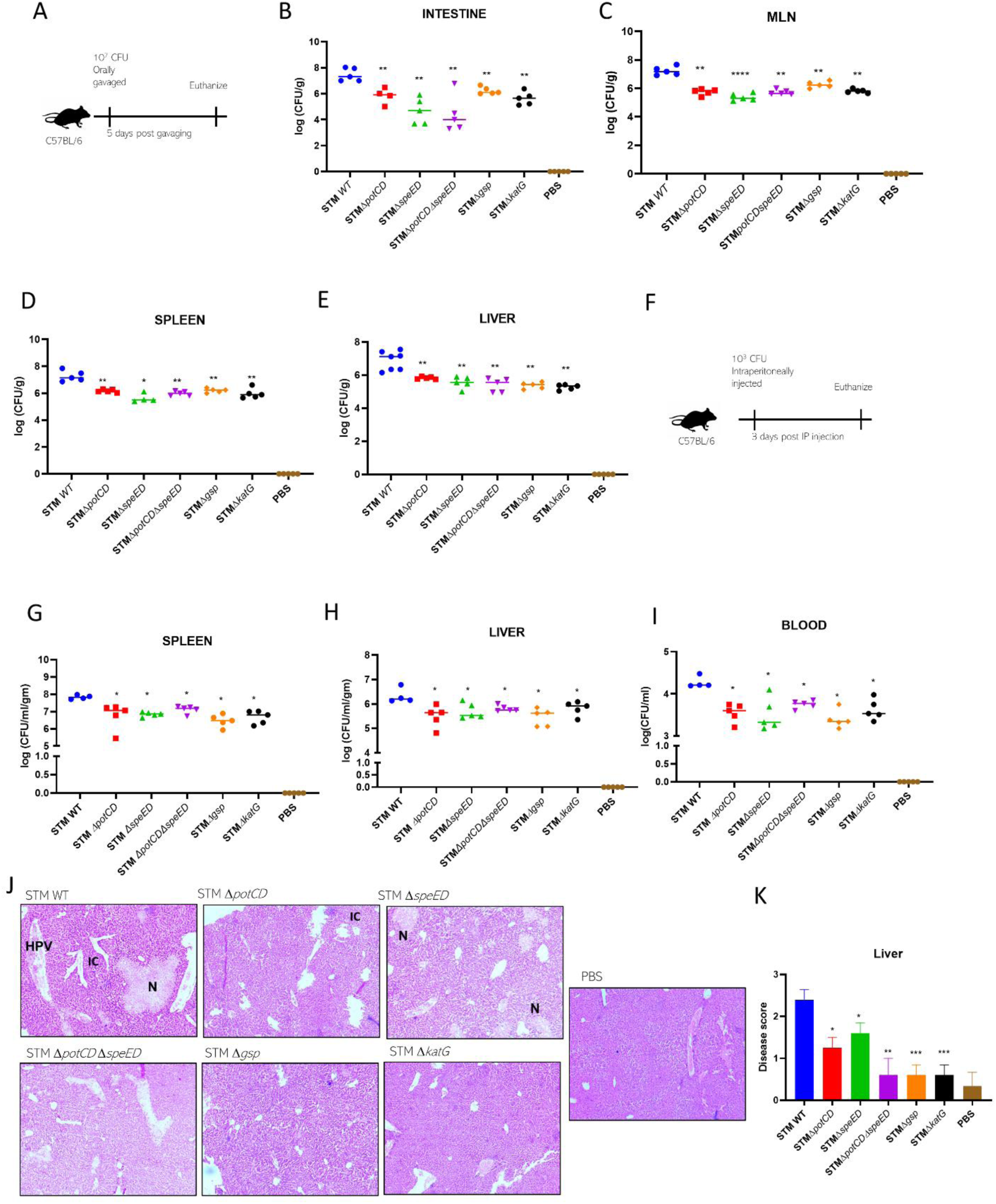
Spermidine is critical for *Salmonella* Typhimurium to colonise the primary and secondary sites of infection in mice. A.The experimental protocol for organ burden in C57BL/6 mice by orally gavaging 10^7^ CFU per mice, B. The organ burden post 5 days of oral gavage in intestine, C. in Mesentric lymph node (MLN), D. in spleen, E. in liver, F. The experimental protocol for organ burden in C57BL/6 mice upon intraperitoneal (I.P) infection with 10^3^ CFU per mice, G. The organ burden 3 days post I.P infection in spleen, H. in liver, I. dissemination in blood, J. The hematoxylin and eosin staining of the sections of liver 3 days post I.P infection of C57Bl/6 mice, K. Disease score from the histopathogical sections of liver, Here, Small necrotic areas (N), Congestion and damage to the endothelial lining of the central vein (C), Congestion of the hepatic portal vein and poor hepatic architecture (HPV), Inflammatory immune cells (IC). The disease score is as :0 for normal pathology, 1 for mild/ minor pathology, 2 for moderate pathology, and 3 for severe pathological changes. Mann Whitney test was used to analyse organ burden in mice; p values ****<0.0001, ***<0.001, **<0.01, *<0.05. Student’s t-test was used to analyze the data; p values ****<0.0001, ***<0.001, **<0.01, *<0.05.

We observe that STM *ΔpotCD*, STM *ΔspeED* show a higher intracellular ROS than STM *Δgsp.* Thus to understand does STM *Δgsp* phenocopy STM *ΔpotCD*, STM *ΔspeED*, we generated double mutants STM *ΔgspΔpotCD* and STM *ΔgspΔspeED.* We find that, the double mutants show a similar reduced fold proliferation and percentage phagocytosis in RAW 264.7 cells (**Fig S6 A-B)**. However, they show an enhanced intracellular ROS upon exposure to hydrogen peroxide (**Fig S6 C)**. Upon co-infection of the double mutants with STM *Δgsp* in C57BL/6 mice, we see that STM *Δgsp* out competes the double mutants in colonizing the liver (**Fig S6 D-E)**. To further dissect the role of spermidine in protection of *Salmonella* Typhimurium from oxidative stress, we infected STM WT, STM *ΔpotCD*, STM *ΔspeED*, STM *ΔpotCDΔspeE,* STM *Δgsp* and STM *ΔkatG* in primary macrophages isolated from the peritoneal lavage of *gp91phox -/-* mice. Gp91Phox is the major subunit of the NOX2 complex, that aids in the catalysis of oxygen to superoxide radical. Interestingly we observe a rescue in the attenuated fold proliferation of STM *ΔpotCD*, STM *ΔspeED*, STM *ΔpotCDΔspeE,* STM *Δgsp* and STM *ΔkatG* in peritoneal macrophages isolated from *gp91phox -/-* mice (**Fig 3 I**). Our results demonstrate the vital role of spermidine in oxidative stress resistance in *Salmonella* Typhimurium.

### Spermidine is critical for *Salmonella* Typhimurium to colonise the primary and secondary sites of infection in mice

*Salmonella* infects the host and breaching the epithelial cells at the Peyer’s patches in the distal ileum, it disseminates to the secondary sites of infection namely the Mesentric Lymph node (MLN), spleen and liver. From the basolateral surface of the epithelial cells at the lamina propria, *Salmonella* is taken by the macrophages and polymorphonuclear cells (PMN). We observed that the spermidine transporter and biosynthesis mutants show attenuated survival in macrophages and under oxidative stress *in vitro.* We were intrigued to study the behaviour of the mutants during *in vivo* colonisation in the mouse model of *Salmonella* infection. We infected C57BL/6 mice by orally gavaging with STM WT, STM *ΔpotCD*, STM *ΔspeED*, STM *ΔpotCDΔspeE,* STM *Δgsp* and STM *ΔkatG* at a CFU of 10^7^ per mice. We noted that STM *ΔpotCD*, STM *ΔspeED*, STM *ΔpotCDΔspeE,* STM *Δgsp* and STM *ΔkatG* showed a significantly lower organ burden in Peyer’s patches, MLN, spleen and liver upon oral gavage (**Fig 4 A-E)**. Our previous study showed that the spermidine transporter and biosynthesis mutants exhibit a poor invasiveness into IECs, which can explain our *in vivo* colonisation upon oral gavage [16]. Oral gavage mimics the physiological route of *Salmonella* infection into its host. Hence, it requires to be able to breach the intestinal barrier successfully. Incapability to invade the IECs in STM *ΔpotCD*, STM *ΔspeED*, STM *ΔpotCDΔspeE* and STM *Δgsp* explains the diminished colonisation in the organs. To dissect the role of spermidine in *in vivo* colonisation we infected C57Bl/6 mice intraperitoneally, by bypassing the entry by breaching epithelial barrier. We observed that upon infecting intraperitoneally, STM *ΔpotCD*, STM *ΔspeED*, STM *ΔpotCDΔspeE,* STM *Δgsp* and STM *ΔkatG* exhibited reduced colonisation in spleen and liver and less dissemination in blood compared to STM WT (**Fig 4 F-I)**. Also, the histopathological sections show significantly less liver tissue damage with STM *ΔpotCD*, STM *ΔspeED*, STM *ΔpotCDΔspeE,* STM *Δgsp* and STM *ΔkatG*, which is validated by disease scoring of the same(**Fig 4 J and K)**. Our results thus show that spermidine aids in the *in vivo* pathogenesis and virulence of *Salmonella* Typhimurium.

### *Salmonella* rewires host polyamine metabolism to potentiate its survival within host macrophages

Most of the intracellular pathogens establish their persistence in the phagocytic cells and are often found to be associated with different populations of the macrophages. Like *Brucella abortus* preferentially resides in the Alternatively activated macrophages (AAM), where it survives and replicates by exploiting the host polyamines. A research group has shown that the metabolism of AAM is shifted to increase polyamine biosynthesis by *Brucella abortus* and thereby promote the bacterial survival [32]. Similarly, *Salmonella* also resides in macrophages, a way to lead chronic infections. We were interested to know whether *Salmonella* exploits the host polyamines and leads to a rewiring of host cell metabolism. To understand the host-pathogen relationship in depth, we assessed the mRNA expression of Ornithine decarboxylase 1 (*mOdc, mouse Ornithine decarboxylase*) which catalyses the rate-determining step of polyamine biosynthesis and Spermidine synthase (*mSrm, mouse Spermidine synthase*) that synthesises spermidine from putrescine by transferring the aminopropyl group from decarboxylated-S-adenosyl methionine in RAW264.7 cells upon infection with STM WT. We observed that mRNA expression of *mOdc1* and *mSrm* were upregulated at 6 and 16 hours post-infection (**Fig 5 A and B**). Further, the mRNA expression of *mOdc1* and *mSrm* were enhanced in the spleen and liver of C57BL/6 mice 5 days post oral gavage with STM WT (**Fig C-F**). Our results show that *Salmonella* Typhimurium upon infection into the host enhances the expression of host polyamine biosynthesis genes. Moreover, we have previously observed that the *Salmonella* Typhimurium that cannot import spermidine cannot survive and proliferate as much as STM WT in macrophages. To further delve into the role of host acquired polyamines, we knocked down *Odc1* in RAW264.7 cells (**Fig S6 A**). Upon knockdown of *mOdc1*, *Salmonella* Typhimurium showed significantly attenuated proliferation in RAW264.7 cells (**Fig 5 G**). However, the knockdown of *mOdc1* did not alter the percentage of phagocytosis by the macrophages (**Fig S7 C)**. Similarly, we knocked down *Srm* in RAW264.7 cells and observed that knock-down of spermidine synthase in the host compromises the ability of STM WT to proliferate and get phagocytosed by the macrophages (**Fig S7 B, D and E)**.

**Figure 5:**
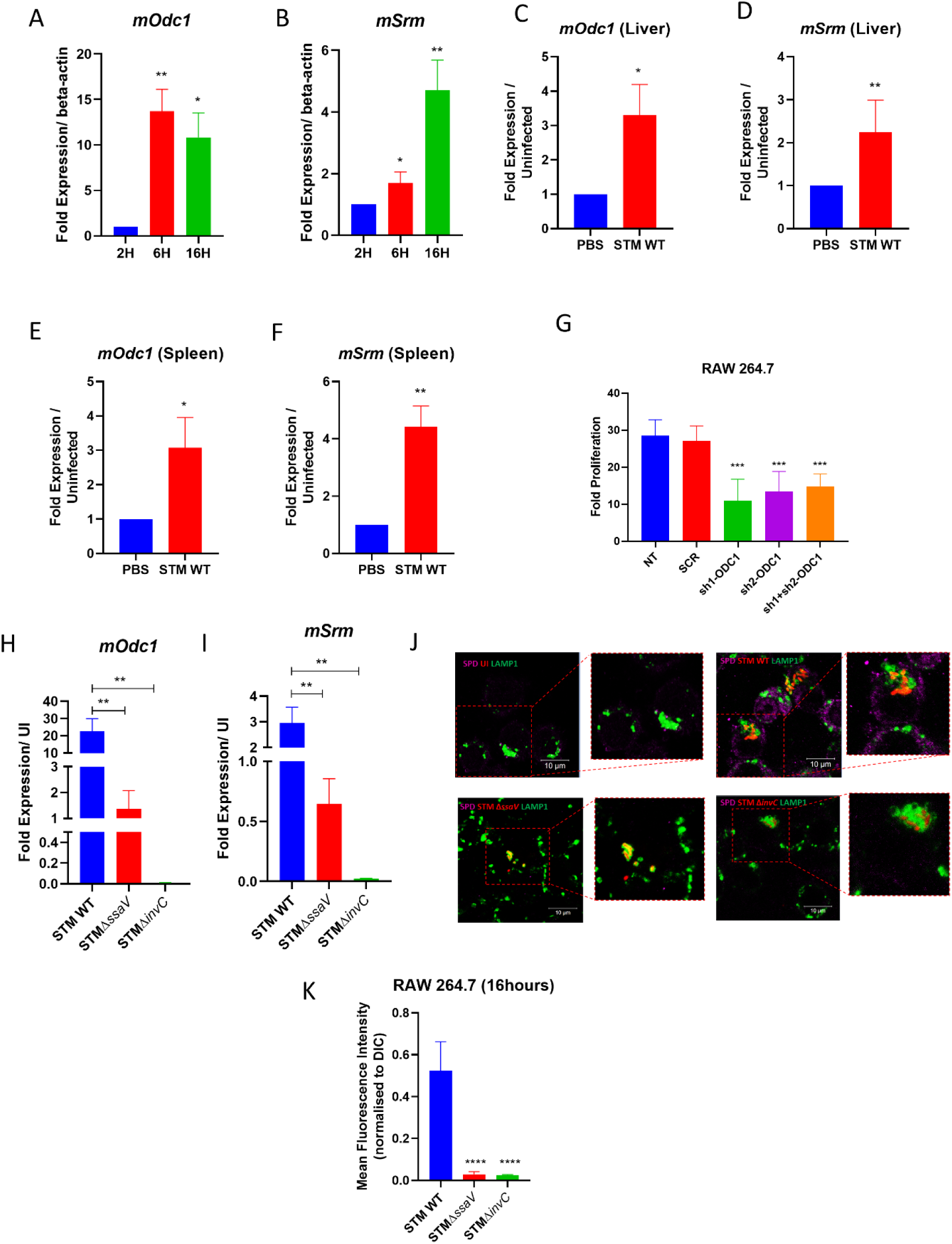
*Salmonella* rewires host polyamine metabolism to potentiate its survival within host macrophages. A.The mRNA expression of *mOdc1* (mouse ornithine decarboxylase) in RAW264.7 cells upon infection with STM WT, B. The mRNA expression of *mSrm* (mouse spermidine synthase) in RAW264.7 cells upon infection with STM WT, C. The mRNA expression of *mOdc1* (mouse ornithine decarboxylase) in liver of C57BL/6 mice 5 days post infection with STM WT by oral gavage, D. The mRNA expression of *mSrm* (mouse spermidine synthase) in liver of C57BL/6 mice 5 days post infection with STM WT by oral gavage,E. The mRNA expression of *mOdc1* (mouse ornithine decarboxylase) in spleen of C57BL/6 mice 5 days post infection with STM WT by oral gavage, F. The mRNA expression of *mSrm* (mouse spermidine synthase) in spleen of C57BL/6 mice 5 days post infection with STM WT by oral gavage, G. The fold proliferation of STM WT in RAW264.7 cells upon transient knockdown of *mOdc1,* H. The percentage phagocytosis of STM WT in RAW264.7 cells upon transient knockdown of *mOdc1.* Here SCR is Scrambled (no target for knock-down), two different targeted shRNA were used for knock-down purposes, Sh1is shRNA-1 for knock-down, Sh2 is shRNA-2 for knock-down, and Sh1+Sh2 indicates where both the shRNAs were used to obtain the knock-down, H. The mRNA expression of *mOdc1* (mouse ornithine decarboxylase) in RAW264.7 cells upon infection with STM WT, STM Δ*ssaV,* and STM Δ*invC*, normalized to the expression in uninfected macrophages, I. The mRNA expression of *mSrm* (mouse spermidine synthase) in RAW264.7 cells upon infection with STM WT, STM Δ*ssaV,* and STM Δ*invC*, normalized to the expression in uninfected macrophages, J. Immunofluorescence imaging to study spermidine in RAW 264.7 cells upon infection with STM WT, STM Δ*invC,* STM Δ*ssaV*, here green is Anti-mouse LAMP1(Alexa fluor 488), Red is pFPV-M-cherry expressing *Salmonella* strains, magenta is anti-Spermidine (Alexa fluor 647) and UI-uninfected. Student’s t-test was used to analyze the data; p values ****<0.0001, ***<0.001, **<0.01, *<0.05.

The question that arises is how does *Salmonella* regulate the host polyamine metabolic pathways? *Salmonella* utilizes *Salmonella* pathogenicity island 1 and 2 encoded effectors for its entry and survival in the specialized host niches [33, 34]. Although SPI-1 is well studied in the initial process of invasion, recent studies suggests that both SPI-1 and SPI-2 effectors are required for SCV maturation and *Salmonella* survival in the host cells [35–37]. Thus to investigate how *Salmonella* modulates host cell polyamine metabolism, we infected RAW264.7 cells with SPI-1 mutant (STM Δ*invC*) and a SPI-2 mutant (STM Δ*ssaV*), both of which lack the ability to import effectors of SPI-1 and SPI-2 respectively. We observed that the mRNA expression of *mOdc1* and *mSrm*, were significantly downregulated in macrophages infected with STM Δ*invC* and STM Δ*ssaV* compared to in STM WT (all normalized to Unifected) (**Fig 5H and I)**. We further determined the intracellular spermidine by immunofluorescence and found that macrophages infected with STM Δ*invC* and STM Δ*ssaV* showed reduced spermidine production compared to in STM WT and uninfected cells (**Fig 5J and K)**. Thus, our data suggests that *Salmonella* utilizes effectors from SPI-1 and SPI-2 to modulate the host cell polyamine metabolic pathways.

### The chemopreventive drug DFMO, reduces *Salmonella* Typhimurium burden in the host by enhancing nitric oxide production

Polyamines are essential molecules in eukarotes, with multiple roles in differentiation, proliferation and development. Many studies have shown that polyamine levels are upregulated in cancer cells and elevated levels of polyamines are associated with breast cancer, neuroblastoma, hepatocellular carcinoma, prostate cancer, lung cancer, colorectal cancer and lukemia [38–42]. D,L-α-difluoromethylornithine (DFMO), an inhibitor of ODC was developed as a potent drug to treat cancer in the year 1970 [43]. DFMO irreversibly binds to the active site of ODC and acts as a suicide inhibitor, thereby reducing polyamine levels and having a cytostatic effect. As a single therapeutic agent it was found to be effective only in neuroblastoma, and cinical trial for other cancer type was unsatisfactory [44, 45]. However, DFMO has been successfully developed as a chemopreventive drug, with FDA approval for treatment of Human African Trypanosomiasis (HAT) [46, 47]. To test whether DFMO can be used as a therapeutic drug against *Salmonella* infection, we treated RAW264.7cells with DFMO during the infection with *Salmonella* Typhimurium, and observed a significant attenuation in the fold proliferation of STM WT (**Fig 6 A)**. Studies have shown that DFMO binds to ODC to prevent further production of putrescine from ornithine and also acts on Arginase1 and reduces the available pool of ornithine for polyamine biosynthesis [48, 49]. This, ensures the flux of arginine to be fed into the nitric oxide synthase (NOS2) pathway and leads to elevated levels of nitric oxide in the cell, which in turn negatively acts on ODC to further block polyamine synthesis [50].We assessed the mRNA expression of *mNos2* upon knockdown of *mOdc1* in RAW264.7 cells followed by infection with STM. We observed an upregulation of *mOdc1* mRNA levels at 6 hours and 16 hours post infection in RAW264.7 cells (**Fig 6 B)**. Further we determined the levels of nitric oxide using a cell permeable dye DAF-2DA, and noted that upon treatment with DFMO there was higher production of nitric oxide (**Fig 6 C and D)**. Taken together our results demonstrate the role of DFMO in diminishing the survival and proliferation of *Salmonella* Typhimurium by blocking Odc1 and enhanced production of nitric oxide in murine macrophages.

**Figure 6:**
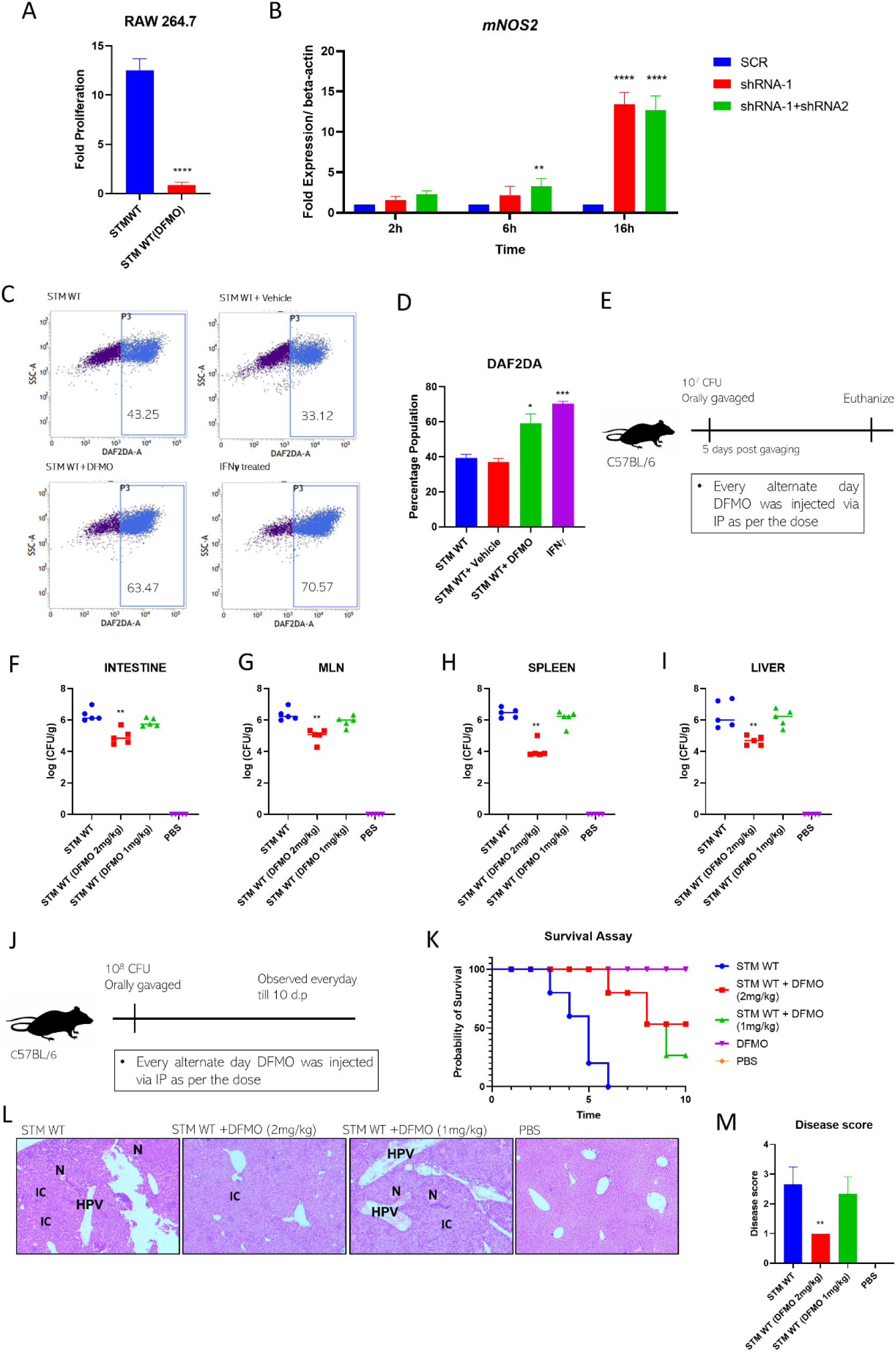
The chemopreventive drug DFMO, reduces *Salmonella* Typhimurium burden in the host by enhancing nitric oxide production. A. The intracellular fold proliferation of STM WT in RAW264.7 cells upon treatment with Difluoromethyl ornithine (DFMO) (5µM), quantified by Gentamicin protection assay, B. The mRNA expression of *mNos2*(mouse Nitric oxide synthase) in RAW264.7 cells, upon transient knockdown of *Odc1* followed by infection with STM WT, the expression is normalized to beta-actin as an internal control, C. The representative scatter plot for DAF2DA dye positive population of RAW264.7 cells, upon transient knockdown of *Odc1* followed by infection with STM WT, the data is determined by flow-cytometry post staining the infected macrophages with the dye, D. The quantification of (C), E. The experimental procedure to study the organ burden of STM WT in C57BL/6 mice upon treatment with DFMO, F-I. The organ burden of STM WT in Intestine, MLN, Spleen and Liver of C57BL/6 mice upon intraperitoneal treatment of DFMO (2mg/kg and 1mg/kg of body weight) as mentioned in (E), J. The Experimental procedure to study the survival of C57BL/6 mice upon infection with STM WT and treatment with DFMO, K.The survival of C57BL/6 mice upon infection with STM WT and upon intraperitoneal treatment of DFMO (2mg/kg and 1mg/kg of body weight) as mentioned in (J), L. Hematoxylin and eosin staining of histopathological sections of liver upon DFMO treatment to C57BL/6 mice. Here, Small necrotic areas (N), Congestion and damage to the endothelial lining of the central vein (C), Congestion of the hepatic portal vein and poor hepatic architecture (HPV), Inflammatory immune cells (IC), M. The disease score index for liver tissue damage upon STM WT infection in C57BL/6 mice with DFMO treatment. The disease score is as :0 for normal pathology, 1 for mild/ minor pathology, 2 for moderate pathology, and 3 for severe pathological changes, Student’s t-test was used to analyze the data; p values ****<0.0001, ***<0.001, **<0.01, *<0.05. Two-way Anova was used to analyze the grouped data; p values ****<0.0001, ***<0.001, **<0.01, *<0.05.

To further address whether DFMO also act as potent anti-*Salmonella* drug in mice model of infection, we infected C57BL/6 mice with 10^7^ CFU per mice by oral gavage. Every alternate day the mice were injected with DFMO in two different doses for two different cohorts (2mg/kg of body weight of mice and 1mg/kg of body weight of mice). Five days post oral gavage we observed that DFMO treatment of 2mg/kg body weight of mice significantly reduced the colonisation of STM WT in intestine, MLN, spleen and liver than in the untreated mice (**Fig 6 E-I)**. Further, using immunofluorescence we observed DFMO significantly reduced the colonisation of STM and and the levels of spermidine in mouse ileum (**Fig S8 A-D).** Moreover, treatment of mice with DFMO at a dose of 2mg/kg body weight of mice, increased the survival of mice upon infection with STM WT (**Fig 6 J and K)**. Also, the weight reduction in mice treated with with DFMO at a dose of 2mg/kg body weight, was less upon infection with STM WT (**Fig S8 E)**. Next, we evaluated the tissue damage upon infection of STM in mice liver. The results show that DFMO treatment of mice at a dose of 2mg/kg of body weight significantly lowered the disease score, suggesting lesser tissue damage compared to untreated (**Fig 6 L and M)**. Thus, DFMO serves as a potential drug to treat *Salmonella* infection in mice by reducing the bacterial burden and tissue damage in mice and enhancing the survival of mice upon infection with STM.

## Discussion

A significant part of the life cycle of *Salmonella* during its pathogenesis involves its stay inside the macrophages. The Gram-negative bacteria come across multiple host environmental stress conditions inside the macrophages [51]. A key mechanism by which the host macrophages try to limit the invading pathogen is by the rapid oxidative burst and ROS production. ROS includes superoxide radicals, hydroxyl radicals, peroxy-nitrites, peroxy-chlorides and hydrogen peroxide. ROS can pass through the bacterial cell wall and act on lipids, proteins and nucleic acids by oxidising them and leading to cellular damage. *Salmonella* is often referred to as a smart pathogen. Over the years, it has evolved to possess multiple strategies to combat the host-derived stresses [52, 53]. To combat the oxidative burst generated NOX2 in macrophages, *Salmonella* possesses multiple antioxidant enzymes such as the catalases KatE and KatG, the superoxide dismutases SodA and SodB, the Alkyl hydroperoxide reductase, the glutaredoxins and thioredoxins and Hrg transcriptional regulator [25]. Polyamines assist in *Salmonella* virulence and aid in stress resistance. However, the mechanism behind the role of polyamines in *Salmonella* stress resistance and virulence remains less appreciated.

Our study identifies spermidine as a stress-responsive regulatory molecule in *Salmonella* Typhimurium. We show spermidine is critical for the survival and proliferation of STM in macrophages and in the presence of oxidative stress *in vitro.* The spermidine transporter and biosynthesis mutants show significant reduced capability to be phagocytosed by the macrophages. Findings from our previous study showed that the intracellular level of spermidine is significantly less in both the spermidine transport and biosynthesis mutants, which further explains the attenuated proliferation of both the mutants in macrophages [16]. The previous findings also showed that in absence of the transport genes in *Salmonella* Typhimurium the synthesis genes are down regulated and vice versa. The absence of spermidine transport and biosynthesis diminishes the mRNA expression of multiple arms of oxidative stress response in *Salmonella.* Multiple studies show the role of polyamines in the regulation of transcription of multiple genes by interacting with DNA in eukaryotes. They bind to the DNA and change conformation as in C-MYC transcription, and in other cases, enhance DNA-protein binding affinities like for Estrogen-response elements [54, 55]. Thus, in *Salmonella* Typhimurium *rpoS, soxR* and *emrR* fall under the “Polyamine modulon”.

We further characterise a novel enzyme GspSA in *Salmonella* Typhimurium, which synthesises a conjugated product of glutathione (GSH) and spermidine called GS-sp. GspSA is critical for *Salmonella* Typhimurium to survive and proliferate in macrophages. Our study shows that the absence of GspSA attenuates the survival of STM under oxidative stress conditions *in vitro,* suggesting a vital role of GspSA in protecting STM from oxidative damage. GS-sp in *E. coli* carries out its function by modifying protein thiol groups under oxidative stress to protect the proteins from getting oxidised and damaged. In *Salmonella* Typhimurium, we further show that the spermidine regulates the *gsp* expression and the subsequent production of GS-sp. We expect that GS-sp generated under oxidative stress conditions and inside the macrophages, likewise interact with cysteine-thiol groups in proteins to shield them from oxidative damage. MK Chattopadhyay (2013) showed that this enzyme (GspSA) is specifically present in two groups of organisms namely bacteria and kineto-plastids respectively and completely absent in other organisms such as human, rats, drosophila etc., among the bacteria group in all *enterobacteria* showed 27-100% homology and >65% identity in around 50% of the species, with *E. coli* [30]. Thus, the absence of GspSA in eukaryotes makes it a potent drug target for treatment of *Salmonella* infections. Thus, spermidine exerts pleiotropic effects in *Salmonella*, by orchestrating the multiple arms of antioxidative response. It strengthens the bacteria to cope with hostile host environments. Our studies in the *in vivo* model of *Salmonella* Typhimurium infection further dissect the role of spermidine in assisting in the pathogenesis by enhancing its virulence properties. Moreover, it is at the nexus of multiple oxidative stress response arms in *Salmonella*, thereby assisting in mounting an antioxidative response to promote its survival in macrophages.

Studies over the past years have given insights into the function of polyamines in the pathogenesis of multiple virulent bacteria. Many human pathogenic bacteria have developed ways to exploit, interfere and manipulate the polyamine metabolism of the host to enhance their fitness within the niche. As in *Shigella* and *Vibrio spp.* polyamines produced by the bacteria are critical in determining virulence [56, 57]. While bacteria such as *Legionella spp*., which does not synthesise polyamines, depend on the host-acquired polyamines for their pathogenesis [58]. Another unique mechanism is observed in *H. pylori* which activates polyamine oxidation, thereby dysregulation the innate-immune response [59]. Also, in response to *H. pylori* infection, the host macrophages increase the arginase activity and ornithine decarboxylase activity to produce polyamines [60]. A recent study showed that SARS-Cov-2 hijacks the host polyamines for its reproduction and infection [61].

Our findings also reveal that *Salmonella* Typhimurium enhances the polyamine production in the host upon infection using its specialized pathogenicity island encoded effectors from SPI-1 and SPI-2, which might be a strategy to hijack the host polyamines for its survival. This further explains another reason for the reduced proliferation observed for the spermidine transport mutant in macrophages. Also, the knockdown of host ornithine decarboxylase attenuates *Salmonella* proliferation in macrophages. The upregulation of Odc1 activity manages to feed the amino acid, L-arginine, into the polyamine biosynthesis and prevent nitric oxide production, which otherwise would be detrimental for the pathogen. Polyamines have been a well-studied as a drug target for treatment of multiple cancers. The oncogene MYC is upregulated in 70 percent of the cancer types. Ornithine decarboxylase (Odc1) rate limiting enzyme of polyamine biosynthesis is transcriptionally activated, directly by MYC, thereby increasing Odc1 levels in cancer [62]. RAS is another essential factor in cell growth and cancer development, and it directly acts on Odc1 to translationally activate Odc1 in cancer cells [63]. Polyamines biosynthesis is also associated with multiple other oncogenes such as AKT1 and mTORC1 [64, 65]. We further show that upon using Odc1 suicide inhibitor, DFMO, *Salmonella* proliferation could be diminished, and it reduces the bacterial burden in mice. The FDA-approved chemopreventive drug, DFMO, for Human African Trypanosomiasis treatment serves to be a potential drug to cure *Salmonella* infection. In the devastating age of increasing antibiotic resistance, such a drug promises to effectively combat deadly pathogens like *Salmonella*.

## Conclusion

A substantial duration of the infection cycle of *Salmonella* involves the macrophages, which present a very hostile environment to the bacteria. However, *Salmonella* is able to survive and proliferate within host macrophages and utilizes it to disseminate to secondary sites of infection. Our study identifies a novel strategy employed by *Salmonella* Typhimurium to counteract oxidative and nitrosative stress within the host. We demonstrate that spermidine is a critical regulatory molecule in *Salmonella* that regulates multiple antioxidative pathways along with a novel antioxidative enzyme (GspSA) in *Salmonella,* to prevent oxidative damage and assist in its virulence in mice. It further rewires host polyamine metabolism in a SPI-1 and SPI-2 dependent manner to prevent nitric oxide production and enhance its survival. In the era of anti-microbial resistance, this study further recognizes an FDA -approved chemopreventive drug, DFMO, which inhibits the host-polyamine metabolism, as a prospective antidote to treat *Salmonella* infection.

## Availability of data and materials

All data generated and analyzed during this study, including the supplementary information files, are incorporated in this article. The data is available from the corresponding author on request.

## Ethics statement

All the animal experiments were approved by the Institutional Animal Ethics Committee, and the Guidelines provided by National Animal Care were strictly followed during all experiments. (Registration No: 435 48/1999/CPCSEA).

## Acknowledgement

Prof. G. Subba Rao from MCB, IISc is duly acknowledged for providing the for knockdown generation. Divisional Mass Spectrometry facility IISc and Mrs. Sunita Joshi for the MS analysis. Departmental Confocal Facility. Departmental Real-Time PCR Facility, Divisional Flowcytometry facility and Central Animal Facility at IISc are duly acknowledged. Mr Sumith and Ms Navya are acknowledged for their help in image acquisition. Dr. Ritika Chatterjee, Mr. Amartya Mukherjee, Mr. Sushovan Bhattacharyya and Ms Bhavya Joshi are also acknowledged for technical help.

## Author contribution statement

AVN and DC conceived the study. AVN and DC designed experiments. AVN and AS performed experiments. RSR analysed for the disease score of tissue samples. AVN, analyzed the data, prepared the figures and wrote the manuscript draft. AVN, AS and DC reviewed and edited the manuscript. DC supervised the work. All the authors read and approved the manuscript.

