## Supplementary Material for "*Salmonella* Typhimurium employ spermidine to exert protection against ROS-mediated cytotoxicity and rewires host polyamine metabolism to ameliorate its survival in macrophages"

Running Head: *Salmonella* mounts an antioxidative response using de-novo and host-acquired spermidine

Keywords: Spermidine, Macrophages, Antioxidative response, Glutathionyl-spermidine synthetase, Difluoromethyl ornithine

### Figure S1

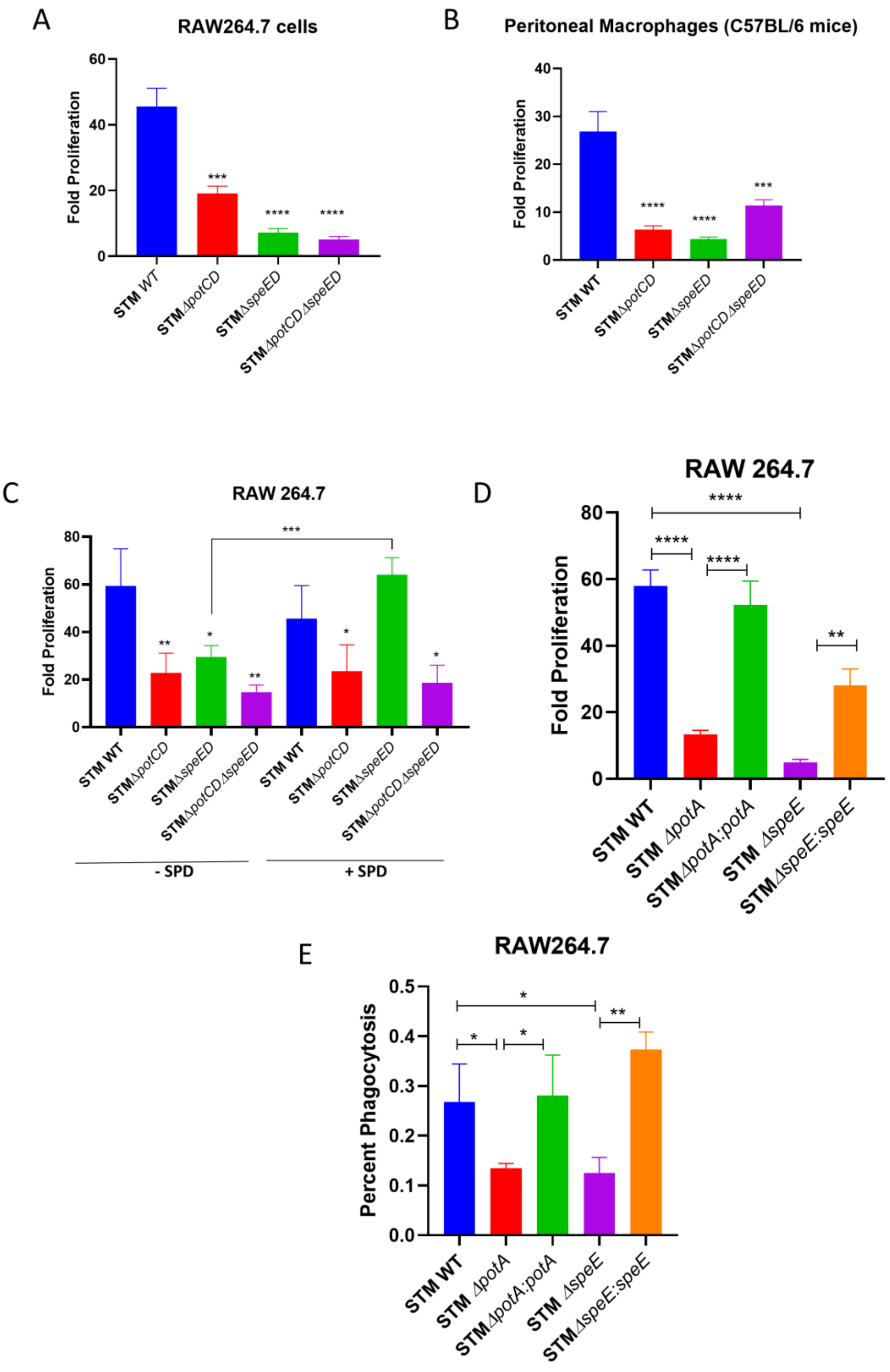
A
RAW264.7 cells
Fold Proliferation
STM WT
STMΔpotCD
STMΔspeED
STMΔpotCDΔspeED
B
Peritoneal Macrophages (C57BL/6 mice)
Fold Proliferation
C
RAW 264.7
Fold Proliferation
- SPD
+ SPD
D
RAW 264.7
Fold Proliferation
STM ΔpotA
STMΔpotA:potA
STM ΔspeE
STMΔspeE:speE
E
RAW264.7
Percent Phagocytosis

Figure S1:

A.The fold proliferation of spermidine mutants in RAW264.7 cells, B. The fold proliferation in primary macrophages isolated from the peritoneal lavage of C57BL/6 mice, C. The fold proliferation of the spermidine mutants grown in media supplemented with 100µM spermidine (SPD) prior to infection, D. The fold proliferation of spermidine mutant complemented strains in RAW264.7 cells, E. The percentage phagocytosis of spermidine mutant complemented strains in RAW264.7 cells Student's t-test was used to analyze the data; p values ****<0.0001, ***<0.001, **<0.01

### Figure S2

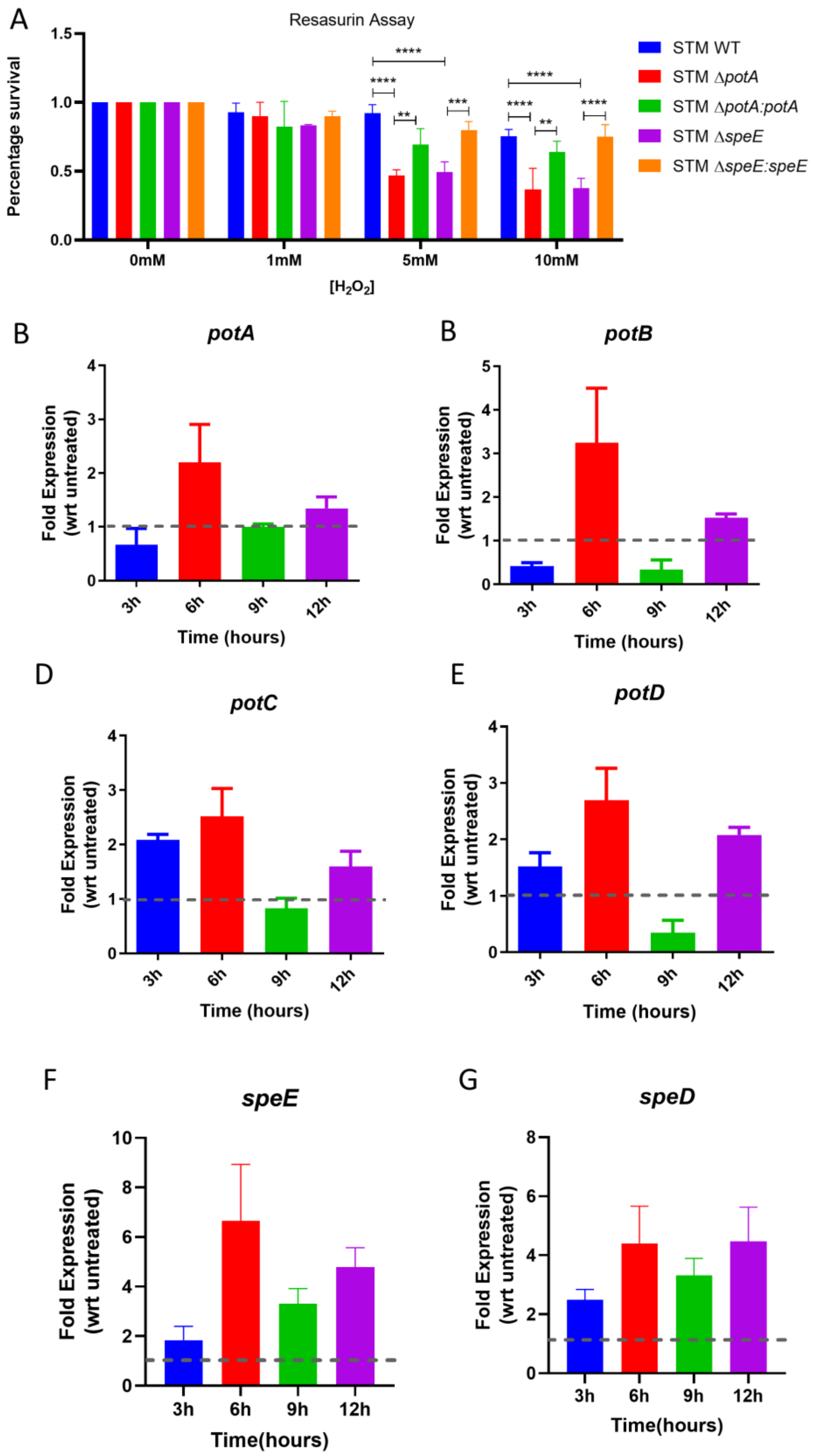
A
Resasurin Assay
Percentage survival
1.5
1.0
0.5
0.0
****
**
***
0mM
1mM
5mM
10mM
[H2O2]
STM WT
STM ΔpotA
STM ΔpotA:potA
STM ΔspeE
STM ΔspeE:speE
B
potA
Fold Expression (wrt untreated)
Time (hours)
3h
6h
9h
12h
B
potB
D
potC
E
potD
F
speE
Time(hours)
G
speD

Figure S2:

A.The hydrogen peroxide sensitivity assay of spermidine mutant complemented strains *in vitro* by resasurin assay, B.The mRNA expression of *potA,* C. *potB,* D. *potC,* E. *potD* in STM WT during its *in vitro* growth in LB media with exposure to oxidative stress (1mM Hydrogen peroxide), Data is normalized to internal control of 16S and represented with respect to the untreated (no hydrogen peroxide), F. The mRNA expression of *speE,* G. *speD* in STM WT during its *in vitro* growth in LB media with exposure to oxidative stress (1mM Hydrogen peroxide), Data is normalized to internal control of 16S and represented with respect to the untreated (no hydrogen peroxide).

### Figure S3

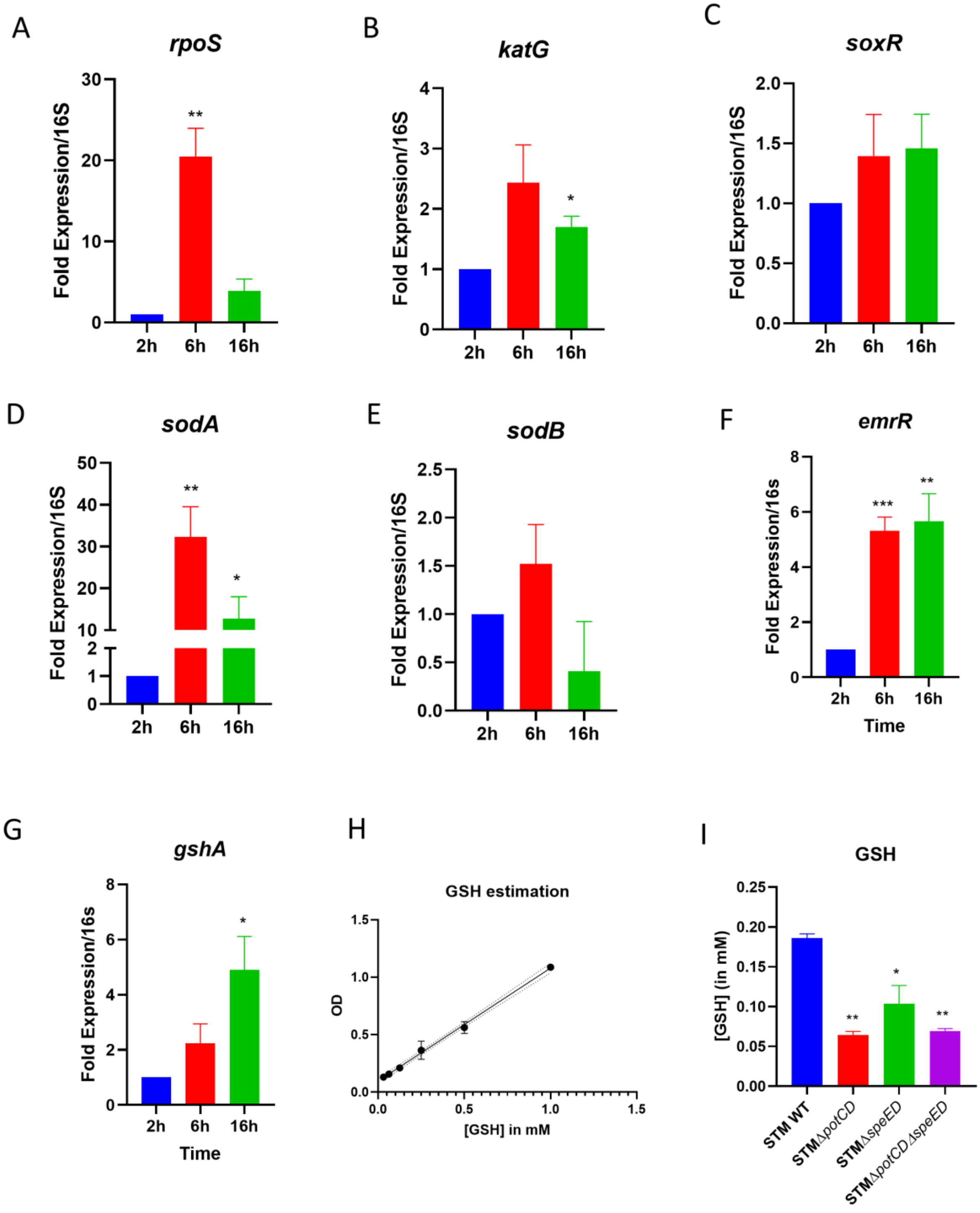
A
rpoS
Fold Expression/16S
30
20
10
0
**
2h
6h
16h
B
katG
Fold Expression/16S
4
3
2
1
0
*
2h
6h
16h
C
soxR
Fold Expression/16S
2.0
1.5
1.0
0.5
0.0
2h
6h
16h
D
sodA
Fold Expression/16S
50
40
30
20
10
2
1
0
**
*
2h
6h
16h
E
sodB
Fold Expression/16S
2.5
2.0
1.5
1.0
0.5
0.0
2h
6h
16h
F
emrR
Fold Expression/16s
8
6
4
2
0
***
**
2h
6h
16h
Time
G
gshA
Fold Expression/16s
8
6
4
2
0
*
2h
6h
16h
Time
H
GSH estimation
OD
1.5
1.0
0.5
0.0
0.0
0.5
1.0
1.5
[GSH] in mM
I
GSH
[GSH] (in mM)
0.25
0.20
0.15
0.10
0.05
0.00
**
*
**
STM WT
STMΔpotCD
STMΔspeED
STMΔpotCDΔspeED

Figure S3:

The mRNA expression in STM WT upon infection into RAW264.7 cells for various antioxidative genes and their transcription factors A. *rpoS*, B. *katG,* C. *soxR,* D. *sodA,* E. *sodB,* F. *emrR,* G. *gshA.* H. Standard curve for Glutathione (GSH) estimation, I. The intracellular GSH concentration determination in spermidine mutants. Student’s t-test was used to analyze the data; p values ****<0.0001, ***<0.001, **<0.01

### Figure S4

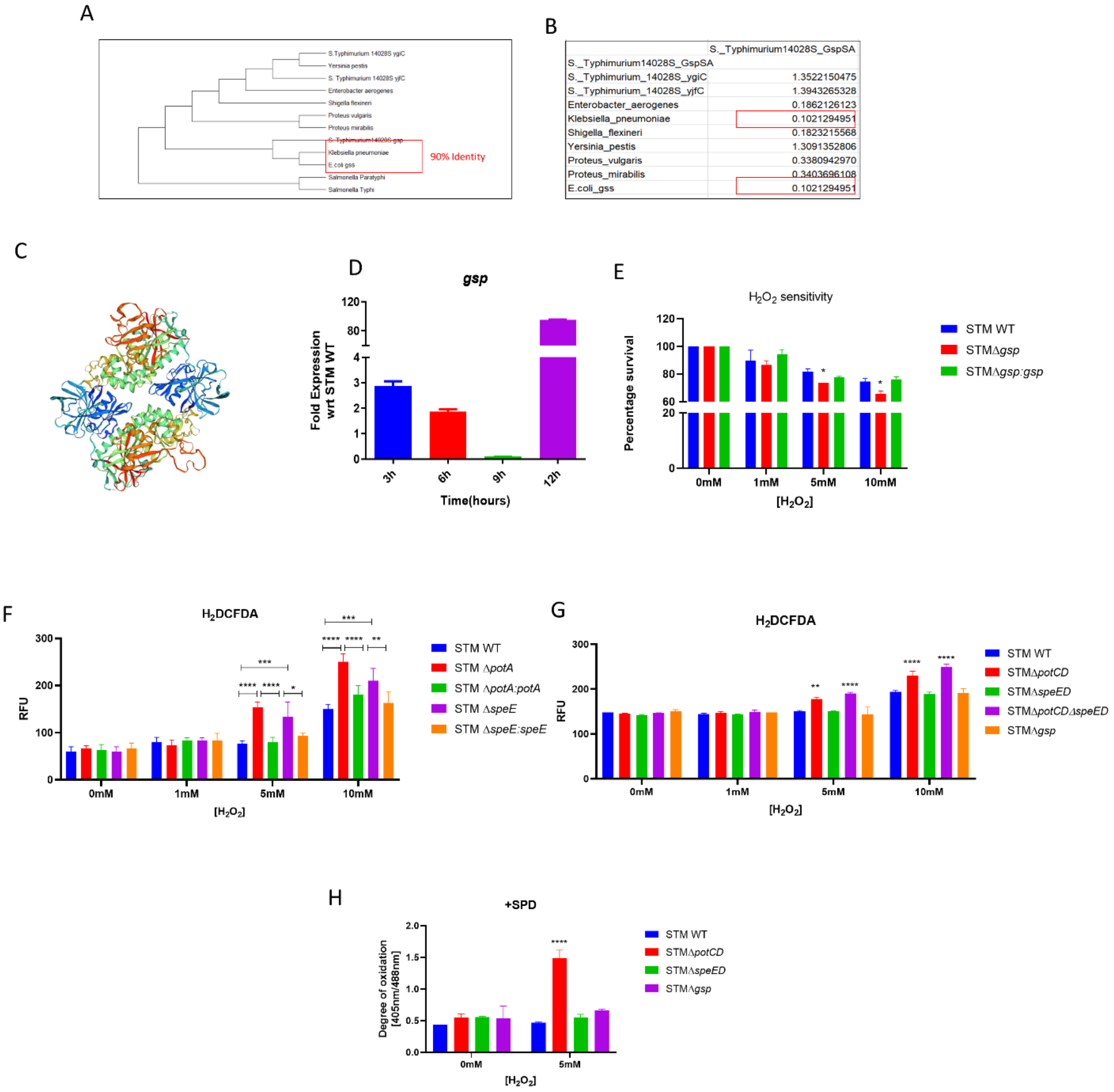


Figure S4:

A.The maximum likelihood tree for GspSA of *Salmonella* Typhimurium 14028S with other members of *Enterobacteriaceae* family, B. The pairwise distance of GspSA in *Salmonella* Typhimurium 14028S from other members of *Enterobacteriaceae* family, C. The prediction of structure using SWISS MODEL , the structure of GspSA in *Salmonella* Typhimurium depicts a homo-dimer with GMQE of 0.93 and QMEANisCo Global of 0.88± 0.05, D. The mRNA expression of *gsp* in STM WT during its *in vitro* growth in LB media upon exposure of 1mM hydrogen peroxide, E. The *in vitro* sensitivity assay of STM WT, STM Δ*gsp* and STM Δ*gsp*:*gsp* in a gradient concentration of hydrogen peroxide, F. The intracellular reactive oxygen species determination using H2DCFDA of spermidine mutant complemented strains, G. The intracellular reactive oxygen species determination using H2DCFDA of all the strains grown with supplementation of 100µM spermidine (SPD), H. The intracellular redox status determination using ro-GFP2 in all the strains grown with supplementation of 100µM spermidine (SPD), Student's t-test was used to analyze the data; p values ****<0.0001, ***<0.001, **<0.01, *<0.05. Two-way ANOVA was used to analyze the grouped data; p values ****<0.0001, ***<0.001, **<0.01, *<0.05.

### Figure S5

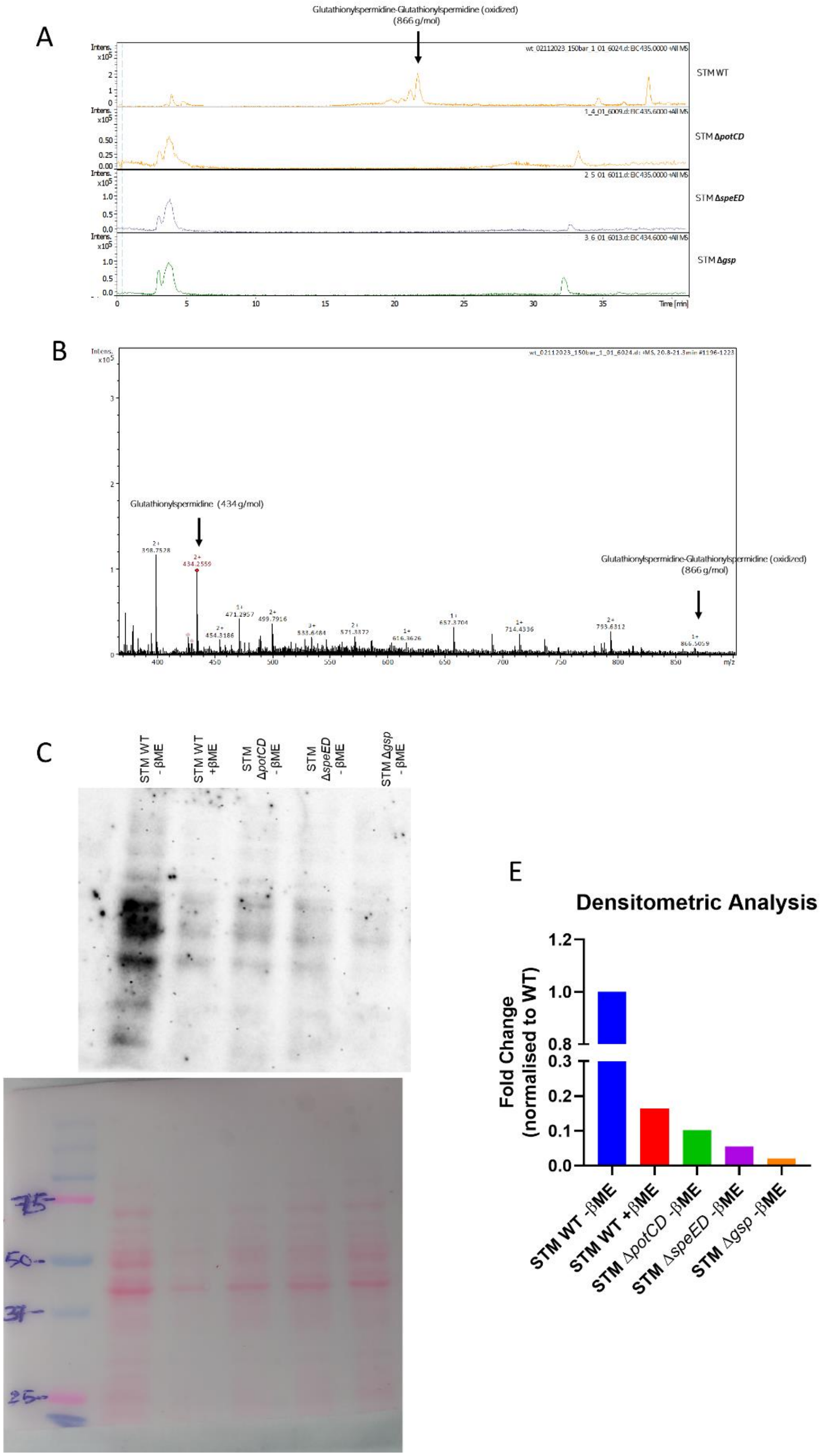
A
Glutathionylspermidine-Glutathionylspermidine (oxidized)
(866 g/mol)
STM WT
STM ΔpotCD
STM ΔspeED
STM Δgsp
Time [min]
B
Glutathionylspermidine (434 g/mol)
Glutathionylspermidine-Glutathionylspermidine (oxidized)
(866 g/mol)
m/z
C
STM WT -βME
STM WT +βME
STM ΔpotCD -βME
STM ΔspeED -βME
STM Δgsp -βME
D
75
50
37
25
E
Densitometric Analysis
Fold Change (normalised to WT)
STM WT -βME
STM WT +βME
STM ΔpotCD -βME
STM ΔspeED -βME
STM Δgsp -βME

Figure S5:

A. The Extracted Ion Chromatogram(EIC) of oxidized Glutathinylspermidine in STM WT, STM Δ*potCD*, STM Δ*speED* and STM Δ*gsp,* the EIC of 435g, B. The mass spectrum showing presence of the oxidized Glutathinylspermidine (866 g/mol) in STM WT sample, C. The immunoblotting for GS-sp-thiolated proteins in STM WT, STM WT+beta mercaptoethanol, STM Δ*potCD*, STM Δ*speED* and STM Δ*gsp* upon exposure to 1mM hydrogen peroxide (anti-spermidine antibody), D. The Ponceau S stain for the above blot in C, E. Densitometric analysis for C (normalized with Ponceau S stain for each lane).

### Figure S6

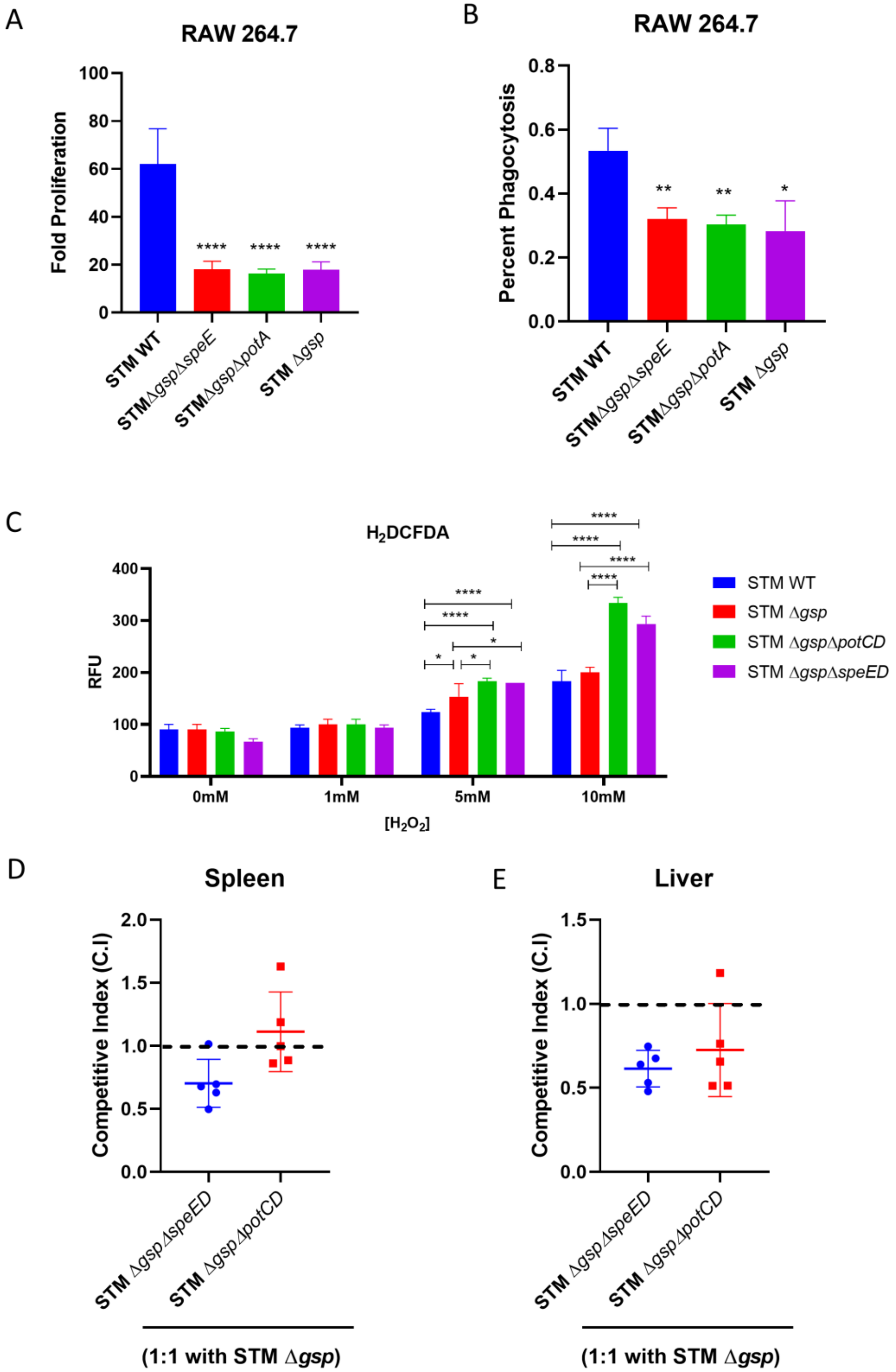
A
RAW 264.7
Fold Proliferation
STM WT
STMΔgspΔspeE
STMΔgspΔpotA
STM Δgsp
B
RAW 264.7
Percent Phagocytosis
C
H2DCFDA
RFU
0mM
1mM
5mM
10mM
[H2O2]
STM WT
STM Δgsp
STM ΔgspΔpotCD
STM ΔgspΔspeED
D
Spleen
Competitive Index (C.I)
STM ΔgspΔspeED
STM ΔgspΔpotCD
(1:1 with STM Δgsp)
E
Liver

Figure S6:

A.The fold proliferation of spermidine and *gsp* double mutant strains in RAW264.7 cells, B. The percentage phagocytosis of spermidine and *gsp* double mutant strains in RAW264.7 cells, C. The intracellular reactive oxygen species determination using H2DCFDA of spermidine and *gsp* double mutant strains, D. The competitive index of spermidine and *gsp* double mutant strains with single *gsp* mutant (1:1 ratio) in Spleen, and E. Liver of C57BL/6 mice Student's t-test was used to analyze the data; p values ****<0.0001, ***<0.001, **<0.01, Mann-Whitney test was used to analyse animal experimental data, p values ****<0.0001, ***<0.001, **<0.01

### Figure S7

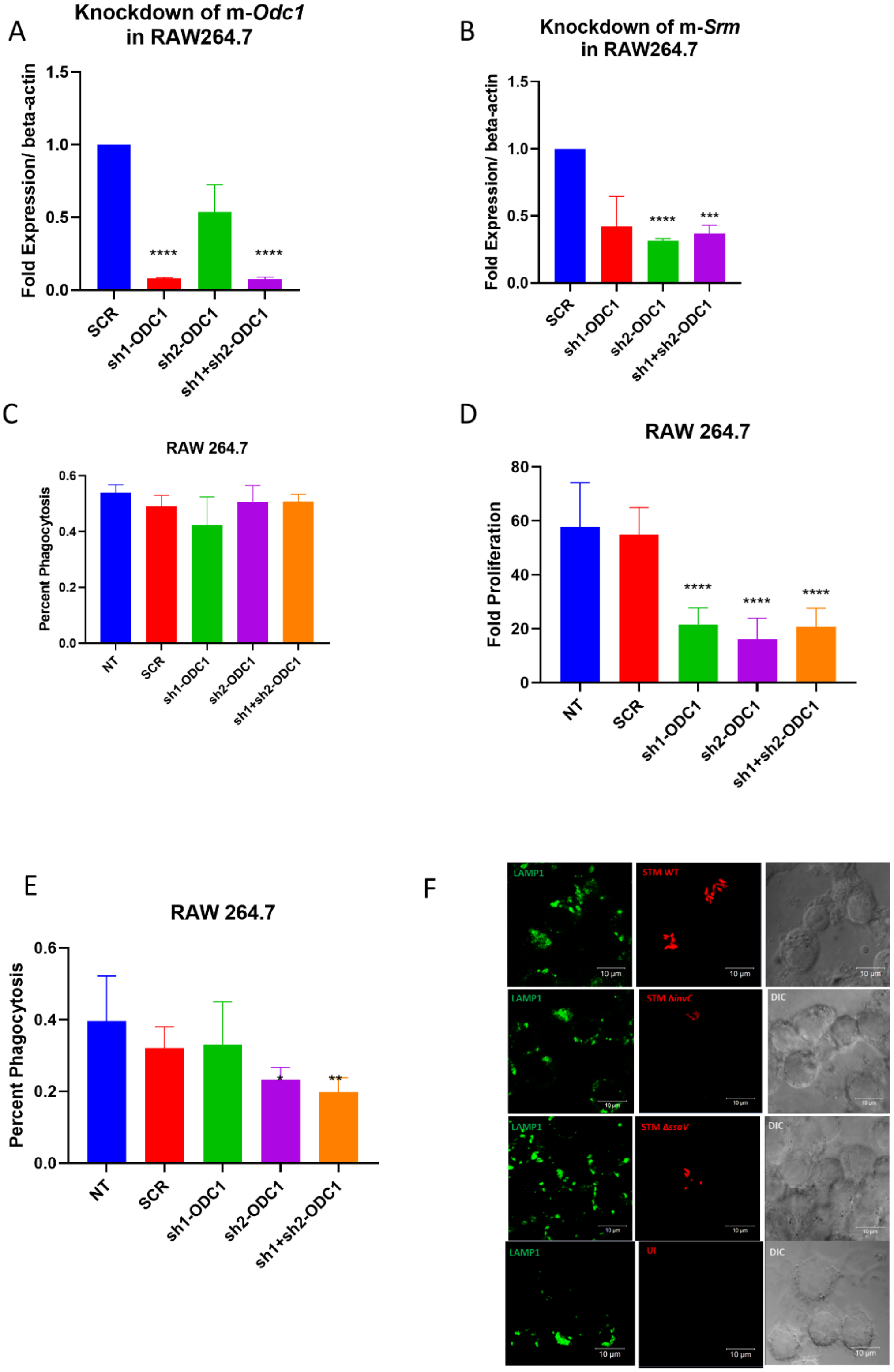

A
Knockdown of m-Odc1 in RAW264.7
Fold Expression/ beta-actin
1.5
1.0
0.5
0.0
****
****
SCR
sh1-ODC1
sh2-ODC1
sh1+sh2-ODC1
B
Knockdown of m-Srm in RAW264.7
Fold Expression/ beta-actin
1.5
1.0
0.5
0.0
****
***
SCR
sh1-ODC1
sh2-ODC1
sh1+sh2-ODC1
C
RAW 264.7
Percent Phagocytosis
0.6
0.4
0.2
0.0
NT
SCR
sh1-ODC1
sh2-ODC1
sh1+sh2-ODC1
D
RAW 264.7
Fold Proliferation
80
60
40
20
0
****
****
****
NT
SCR
sh1-ODC1
sh2-ODC1
sh1+sh2-ODC1
E
RAW 264.7
Percent Phagocytosis
0.6
0.4
0.2
0.0
*
**
NT
SCR
sh1-ODC1
sh2-ODC1
sh1+sh2-ODC1
F
LAMP1
STM WT
STM ΔinvC
STM ΔssaV
UI
DIC
10 μm

Figure S7:

A.Confirmation of knock-down of Odc1 in RAW264.7 cells using qRT-PCR, B. Confirmation of knock-down of Srm in RAW264.7 cells using qRT-PCR, C. The percentage phagocytosis of STM WT in RAW264.7 cells upon transient knock-down of Odc1, D. The fold proliferation of STM WT in RAW264.7 cells upon transient knock-down of Srm, E. The percentage phagocytosis of STM WT in RAW264.7 cells upon transient knock-down of Srm. Here SCR is Scrambled (no target for knock-down), two different targeted shRNA were used for knock-down purposes, Sh1is shRNA-1 for knock-down, Sh2 is shRNA-2 for knock-down, and Sh1+Sh2 indicates where both the shRNAs were used to obtain the knock-down, F. Immunofluorescence imaging to study the spermidine in RAW 264.7 cells upon infection with STM WT, STM Δ*invC,* STM Δ*ssaV*, here green is Anti-mouse LAMP1(Alexa fluor 488), Red is pFPV-M-cherry expressing *Salmonella* strains, magenta is anti-Spermidine (Alexa fluor 647) and UI- uninfected Student's t-test was used to analyze the data; p values ****<0.0001, ***<0.001, **<0.01, *<0.05.

### Figure S8

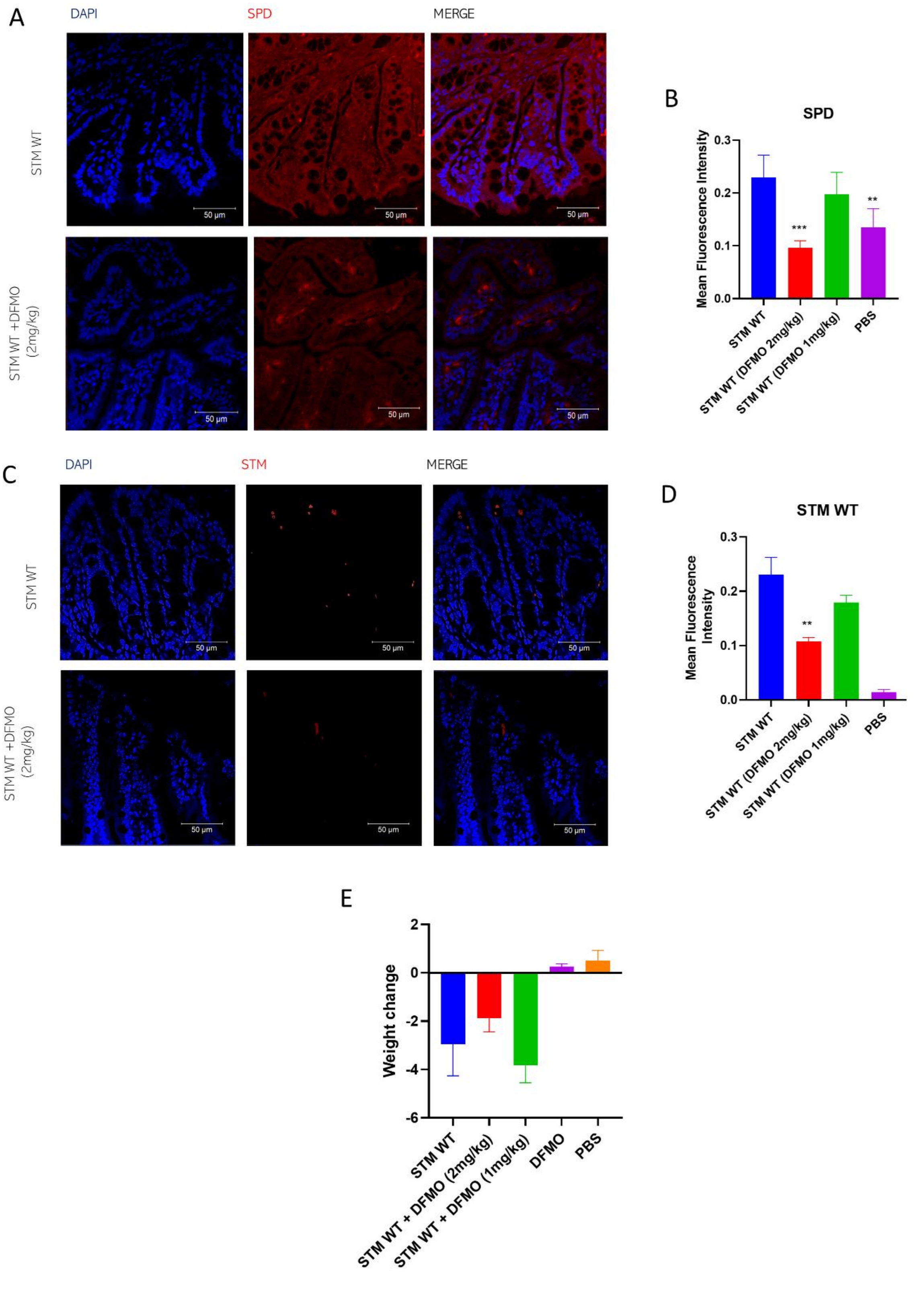
A
DAPI
SPD
MERGE
STM WT
STM WT +DFMO (2mg/kg)
50 μm
B
SPD
Mean Fluorescence Intensity
0.3
0.2
0.1
0.0
***
**
STM WT
STM WT (DFMO 2mg/kg)
STM WT (DFMO 1mg/kg)
PBS
C
DAPI
STM
MERGE
STM WT
STM WT +DFMO (2mg/kg)
50 μm
D
STM WT
Mean Fluorescence Intensity
0.3
0.2
0.1
0.0
**
STM WT
STM WT (DFMO 2mg/kg)
STM WT (DFMO 1mg/kg)
PBS
E
Weight change
2
0
-2
-4
-6
STM WT
STM WT + DFMO (2mg/kg)
STM WT + DFMO (1mg/kg)
DFMO
PBS

Figure S8:

A. Immunofluorescence imaging of mice ileum showing the inhibitory effect of DFMO on polyamine biosynthesis in mice. Here DAPI(blue) stains the intestinal cells and Anti-spermidine antibody is used to stain for spermidine (Cy3 tagged secondary antibody (red)). Representative images are shown, B.The quantification of (A) by Mean fluorescence intensity, C. Immunofluorescence imaging of mice ileum showing the colonisation of STM in mice upon DFMO treatment. Here DAPI(blue) stains the intestinal cells, and Anti-*Salmonella*-LPS antibody is used to stain for STM (Cy3 tagged secondary antibody (red)). Representative images are shown, D. The quantification of (C) by Mean fluorescence intensity, E. The weight change (reduction) of mice upon infection with STM WT and followed the intraperitoneal treatment of DFMO (2mg/kg and 1mg/kg of body weight) every alternate day. Data acquired 5 days post infection. Unpaired Student's t-test was used to analyze the data ; p values ****<0.0001, ***<0.001, **<0.01, *<0.05.

### Supplementary Table

#### *Salmonella* Typhimurium employ spermidine to exert protection against ROS-mediated cytotoxicity and rewires host polyamine metabolism to ameliorate its survival in macrophages

Abhilash Vijay Nair[a], Anmol Singh[a], R. S. Rajmani[b] , Dipshikha Chakravortty[a,c]#

[a]Department of Microbiology and Cell Biology, Division of Biological Sciences, Indian Institute of Science, Bengaluru, India

[b]Molecular Biophysiscs Unit, Indian Institute of Science, Bangalore, India

[c]Adjunct Faculty, School of Biology, Indian Institute of Science Education and Research, Thiruvananthapuram

Running Head: *Salmonella* mounts an antioxidative response using de-novo and host-acquired spermidine

Keywords: Spermidine, Macrophages, Antioxidative response, Glutathionyl-spermidine synthetase, Difluoromethyl ornithine

#### S-Table 1: Bacterial Strains

| | | |
|---|---|---|
| A | STM WT (14028s) | No antibiotic |
| B | STM *ΔpotCD* | Generated in background of (A), Kanmycin resistance (From Previous study) |
| C | STM *ΔspeED* | Generated in background of (A), Chloramphenicol resistance (From Previous study) |
| D | STM *ΔpotCDΔspeED* | Generated in background of (A), Kanamycin and Chloramphenicol resistance (From Previous study) |
| E | STM *ΔfliC* | Generated in background of (A), Kanmycin resistance ( Lab stock) |
| F | STM *Δgsp* | Generated in background of (A), Kanmycin resistance (This study) |
| G | STM *Δgsp:gsp* | Generated in background of (A), Kanmycin resistance and complemented through pQE60 plasmid (Ampicillin resistance) (This study) |
| H | STM *ΔkatG* | Generated in background of (A), Kanmycin resistance (Lab stock) |
| I | STM *ΔinvC* | Generated in background of (A), Kanmycin resistance (Lab stock) |
| J | STM *ΔssaV* | Generated in background of (A), Kanmycin resistance (Lab stock) |

#### S-Table 2: Primers

| Primer | Sequence |
|---|---|
| *gsp* Knockout primer FP | 5’-TCTTGTTTTGAGGTAAGTGCATGAGCAAAGGAACCACCAGCATATGAATATCCTCCTTAG -3’ |
| *gsp* Knockout primer RP | 5’-AATGACCCCGTTACGCTTTGATGACGATTAACGGTTCAATGTGTAGGCTGGAGCTGCTTC -3’ |
| *gsp* Knockout confirmation primer FP | 5’-TCTGCCATATTGACCTGCCT -3’ |
| *gsp* Knockout confirmation primer RP | 5’-AACAGGCATGACCTAAATCC -3’ |
| *potA* Expression primer FP | 5’-AAGGGCTTATCGGCTACGTG-3’ |
| *potA* Expression primer RP | 5’-CATCAGCCAGTACGACCTCC-3’ |
| *potB* Expression primer FP | 5’-CCTTTTGCCTGGTTTCTGGC-3’ |
| *potB* Expression primer RP | 5’-CGGTGTATCAATCACGCCCA-3’ |
| *potC* Expression primer FP | 5’-GAATGCTGGAAGCCGCAAAA -3’ |
| *potC* Expression primer RP | 5’-TTCTGGTGAAACGCCGACTT -3’ |
| *potD* Expression primer FP | 5’-TCCGAAAGAGATCGAAGCGG -3’ |
| *potD* Expression primer RP | 5’-TTTTTCGCGTTTGCCGGAAT -3’ |

| | |
|---|---|
| *speE* Expression primer FP | 5'-GCGGCGATTCCGACCTATTA -3' |
| *speE* Expression primer RP | 5'-CGGACAGTGCGTCATGTAGA -3' |
| *speD* Expression primer FP | 5'-CTACGCCAAAACCGCAGAAG -3' |
| *speD* Expression primer RP | 5'-GGCTCTTCGCTCACCAGAAT-3' |
| *16S* Expression primer FP | 5'- GTGAGGTAACGGCTCACCAA-3' |
| *16S* Expression primer RP | 5'- TAACCGCAACACCTTCCTCC -3' |
| *hilA* Expression primer FP | 5'-CGCCGGCGAGATTGTGAGTA -3' |
| *hilA* Expression primer RP | 5'-TGGCGGAGACACCACTACGA -3' |
| *sipA* Expression primer FP | 5'-TTCGGATGAAGCGTTGGTCA -3' |
| *sipA* Expression primer RP | 5'-GTCATTCGCGTGTGGATTCG -3' |
| *sopD* Expression primer FP | 5'-TGGCGGAGACACCACTACGA -3' |
| *sopD* Expression primer RP | 5'-TGGCGGAGACACCACTACGA -3' |
| *sodA* Expression primer FP | 5'- GCGTGTTCCCACACATCAAC -3' |
| *sodA* Expression primer RP | 5'- GCTATCGCGGCATCTTTTGG -3' |
| *sodB* Expression primer FP | 5'- CCTGCCGGTTGAAGAACTGA -3' |
| *sodB* Expression primer RP | 5'-ACCGAAGTCACGCTCGATAG -3' |
| *katG* Expression primer FP | 5'-TGGCGGAGACACCACTACGA -3' |
| *katG* Expression primer RP | 5'- CACGGTCTCTTCGTCGTTCA -3' |
| *gsp* Expression primer FP | 5'-TTTTCTGCGTGGCGAACTTG-3' |
| *gsp* Expression primer RP | 5'-GTGTTTGAACCGCTGTGGAC -3' |
| *gshA* Expression primer FP | 5'- GCCGCTTACCGTCTTTTTCC -3' |
| *gshA* Expression primer RP | 5'- AGTCGCAAAGCAATCTCGGA -3' |
| *emrR* Expression primer FP | 5'- TGGATTGAGCGTCGTGAGAG -3' |
| *emrR* Expression primer RP | 5'- CGCGTCAGGAGTTTACGAGT -3' |

| | |
|---|---|
| *Mouse Odc1* Expression primer FP | 5’-ATCTCCAGGTTCCCTGTCACAT -3’ |
| *Mouse Odc1* Expression primer RP | 5’-AATGTCTCCGAGGTCCGCAA-3’ |
| *Mouse Srm* Expression primer FP | 5’-TCCTCGTCTTCCGCAGTAAA -3’ |
| *Mouse Srm* primer RP | 5’-GTGCTTCACCACTTCCCGTA -3’ |
| *Mouse beta-actin* Expression primer FP | 5’-CAGCAAGCAGGAGTACGATG-3’ |
| *Mouse beta-actin* Expression primer RP | 5’-GCAGCTCAGTAACAGTCCG -3’ |

#### S-Table 3: shRNA sequences

| **shRNA used for knockdown** | | |
|---|---|---|
| **Name** | **TRC ID** | **Selected shRNA** |
| Mouse ODC1 A7 | TRCN0000344821 | CCGGGCCG**ACGATCTACTATGTGATCTC** GAGATCACATAGTAGATCGTCGGCTTTTT G |
| Mouse ODC1 A8 | TRCN0000333415 | CCGGACGG**GCGAAAGAGCTAAATATCTC** GAGATATTTAGCTCTTTCGCCCGTTTTTG |
| Mouse SRM E9 | TRCN0000290713 | CCGGAGTC**TCTTCAAGGAGTCCTATCTC** GAGATAGGACTCCTTGAAGAGACTTTTTT G |
| Mouse SRM F1 | TRCN0000290784 | CCGGCATC**CAAGTCTCCAAGAAGTTCTC** GAGAACTTCTTGGAGACTTGGATGTTTTT G |
